# Chronic mitochondrial fragmentation elicits a neuroprotective Warburg-like effect in *Drosophila* neurons

**DOI:** 10.1101/2024.08.30.610549

**Authors:** Shlesha Richhariya, Daniel Shin, Matthias Schlichting, Michael Rosbash

## Abstract

Mitochondrial fission and fusion are dynamic and important cellular processes, but the roles of these two very different mitochondrial forms – predominantly spherical and tubular - are not well-characterized in neurons of animals and especially in aging neurons. This is important because neurons are long-lived and mitochondrial dynamics is associated with neurodegenerative diseases. We used here an efficient cell type-specific CRISPR approach to knockout key fission-fusion genes and disrupt mitochondrial dynamics within the inessential clock neurons of *Drosophila*. Surprisingly, fusion is much more important than fission for maintaining long-term neuronal function. Neurons survive chronic mitochondrial fragmentation due to loss of fusion by triggering a cancer-like transcriptomic response. This Warburg effect includes ATF4-mediated upregulation of the aerobic glycolysis gene *Lactate dehydrogenase* (*Ldh*), and LDH is essential to prevent neurodegeneration of neurons deficient in the fusion gene *Opa1*. These results and others provide insights into the intersection of neuronal metabolism, aging and neurodegeneration.

## Introduction

Mitochondria are organelles present in almost all eukaryotic cells. The primary function of mitochondria is to produce cellular energy in the form of ATP through oxidative phosphorylation, but they also participate in several other processes such as calcium signaling and regulating the response to stress (Shen et al., 2022). Mitochondria are dynamic, as they constantly undergo fission and fusion in response to cellular demands of energy and stress (Chan, 2012; Youle and van der Bliek, 2012). Fission generates fragmented mitochondria and involves mitochondrial division, in part to remove damaged mitochondria or mitochondrial DNA and is mainly mediated by the protein Drp1. Mitochondrial fusion generates larger and more tubular mitochondria and involves the joining of two or more mitochondria in response to stress or increased energy demand. Opa1 and mitofusins (Mfn1 and Mfn2 in mammals) regulate the fusion of the inner and outer mitochondrial membranes, respectively (van der Bliek et al., 2013).

Fission and fusion are essential processes as loss of either one is lethal (Wakabayashi et al., 2009) or results in severe disease. Loss of Opa1 results in Optic atrophy (Alexander et al., 2000; Delettre et al., 2000), whereas loss of mitofusins causes Charcot Marie Tooth disease (Züchner et al., 2004). Although fragmented mitochondrial morphology is predominant in and often interacts with proteins associated with neurodegeneration (Cho et al., 2009; Nakamura et al., 2011; Song et al., 2011; Wang et al., 2009), fission can also be critical in some diseased conditions (Shields et al., 2021). Importantly however, the contribution of fragmented mitochondria or the contribution of reduced levels of fused mitochondria, to disease pathology is not generally understood, especially in aging neurons *in vivo*. In addition, it is important to study the role of mitochondrial dynamics in aging neurons without disease as they are long-lived cells with unique energy demands (Trigo et al., 2022).

*Drosophila* neurons are good models as key mammalian genes regulating mitochondrial dynamics are conserved. *Drp1, Opa1* and *Marf* are the single *Drosophila* orthologs of mammalian *Drp1, Opa1* and mitofusins, respectively (Sandoval et al., 2014; Verstreken et al., 2005; Yarosh et al., 2008). Several neurodegenerative diseases have also been successfully modeled in flies (McGurk et al., 2015). In addition, the short *Drosophila* lifespan of ∼2-3 months is conducive to aging studies. Most importantly, *Drosophila* offers unparalleled genetic access, especially with cell type-specific CRISPR-based methods for disrupting gene function in neurons of interest (Port et al., 2020; Schlichting et al., 2019).

Mitochondrial fission and fusion are also essential processes in flies, and robust inhibition of either process in large numbers of neurons is lethal (Trevisan et al., 2018; Verstreken et al., 2005). However, this is not the case in the ∼150 *Drosophila* clock neurons. They are inessential for survival under lab conditions, and CRISPR-knockouts work well in these neurons (Richhariya et al., 2023; Schlichting et al., 2019). Moreover, the morphology of these neurons is well-characterized, and they require a functional molecular clock as well as neuronal activity to drive robust, rhythmic circadian behavior (Ahmad et al., 2021; Schlichting et al., 2019). This behavioral readout is facile and sensitive, so small as well as large effects on clock neuron function can be easily assayed. These neurons can also be purified for transcriptomic analysis (Ma et al., 2021). All of these features contribute to making these neurons and their resulting behavioral output an attractive platform for the study of essential processes like mitochondrial fission and fusion, including the effect of aging. The results identified molecular signatures of neurons lacking key fission-fusion genes as well as their role in neurodegeneration via a CRISPR-based double genetic interaction approach.

## Results

### Cell type-specific CRISPR blocks mitochondrial fission-fusion in clock neurons

To test the role of mitochondrial fission and fusion in neurons, we employed a cell type-specific CRISPR strategy (Fig. 1A), which was previously found to be very efficient in disrupting gene function (Port et al., 2020; Schlichting et al., 2019). We generated to this end, UAS-3X-gRNA lines targeting genes critical for mitochondrial fission (*Drp1*) and fusion (*Opa1, Marf*). When these lines were expressed along with Cas9 in all neurons using either the *elav-* or the *nSyb-Gal4* drivers, there were no viable adults. However, when these same UAS-3X-gRNA were expressed along with Cas9 in the clock neurons using the *CLK856-Gal4* driver (henceforth referred to as *Drp1-mut*, *Opa1-mut* and *Marf-mut* respectively), not only were normal numbers of adult flies obtained, but their lifespans were also unchanged from controls (Fig. S1). We next checked if the mitochondrial morphology labeled by *UAS-mitoGFP* in clock neurons was altered as expected, i.e., hyperfused in the fission mutant and hyperfragmented in the fusion mutant clock neurons (Fig. 1B). Indeed, mitochondria in large PDF clock neuron cell bodies of young (7-10 day old) *Drp1-mut* flies appear as large aggregates compared to the smaller tubular structures in the Cas9 control flies. Conversely, the *Opa1-mut* or *Marf-mut* neuron cell body mitochondria appeared fragmented (Fig 1C). There were similar effects on mitochondrial morphology in other clock neurons such as the LNds (Fig. S2).

**Fig. 1:**
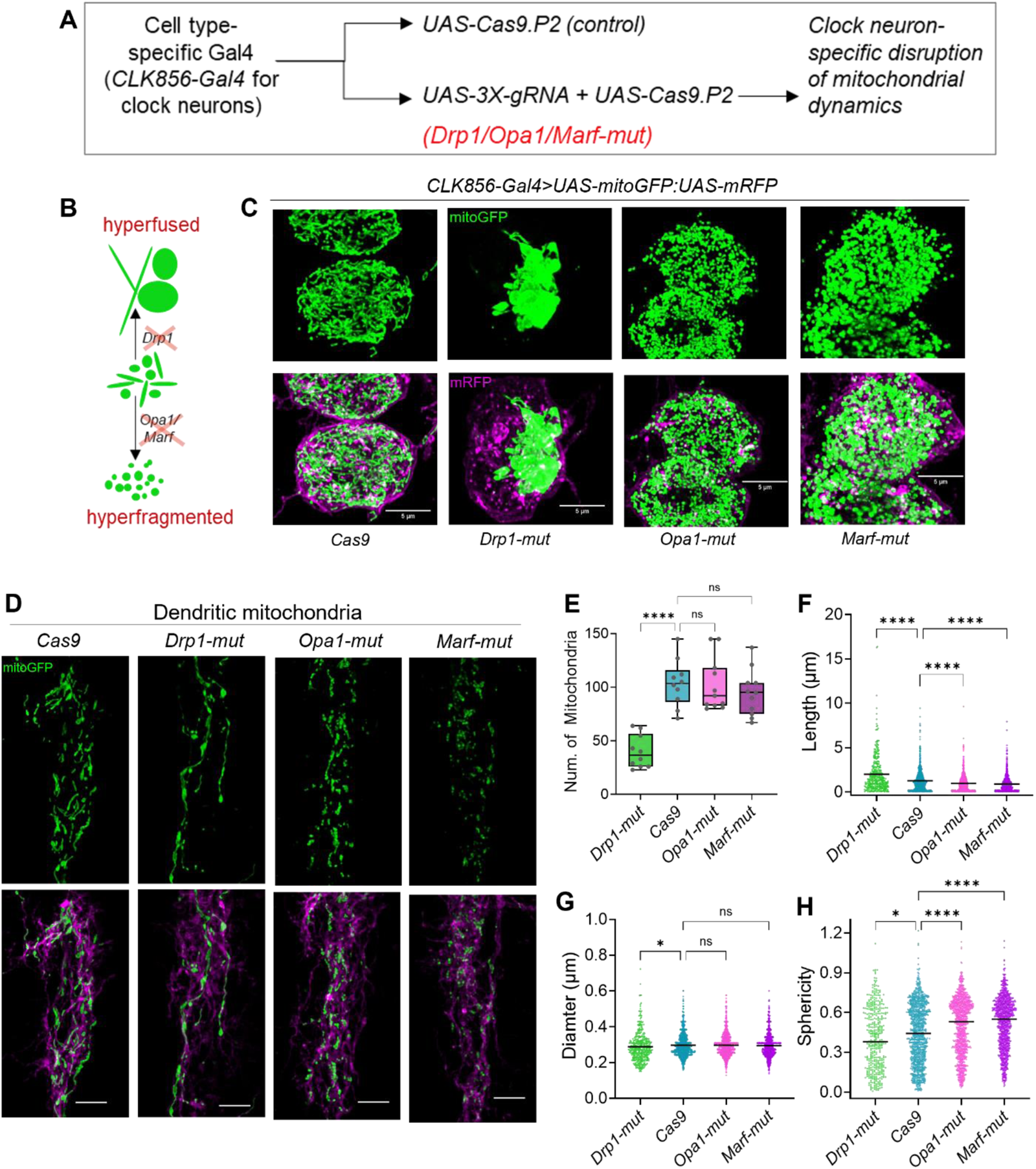
Cell type-specific CRISPR effectively clamps mitochondrial dynamics. **A.** Schematic representation of experimental plan to generate clock neuron-specific mutations in genes regulating mitochondrial dynamics. **B**. Expected mitochondrial morphology from the perturbations – loss of fission gene *Drp1* would lead to hyperfused mitochondria and loss of either fusion genes *Opa1* or *Marf* would lead to fragmented mitochondria. **C**. Representative images of mitochondria labeled with *UAS-mitoGFP* and UAS-*mRFP* in the large PDF neurons, a subset of clock neurons marked by *CLK856-Gal4* with indicated mutations. Scale bars represent 5µm. **D**. Representative images of mitochondria in dendrites of the large-PDF neurons with indicated mutations. **E.** Quantification of number of mitochondria per dendrite. **F-H.** Length, diameter and sphericity of individual dendritic mitochondria. n ≥ 10 dendrites per genotype. Kruskal-Wallis ANOVA with a post hoc Dunn’s test was used to test statistical significance between Cas9-control and the mutants, adjusted p-value >0.05 is denoted as not significant (n.s.); *p<0.05; ****p<0.0001.

We also examined the mitochondrial morphology within the axons and dendrites of the large PDF neurons (Fig. S3A). Consistent with published reports (Rangaraju et al., 2019), control dendritic mitochondria were longer and less spherical than axonal ones (Fig. 1D, S3B). The dendritic mitochondria in *Drp1-mut* neurons were fewer in number, longer, less broad and less spherical, whereas they were shorter and more spherical in both fusion mutants (Fig. 1D-H). Similarly, axonal mitochondria of the fission mutant were fewer in number, longer, more narrow and less spherical. Axonal mitochondria of the fusion mutants were significantly smaller albeit not more spherical than controls (Fig. S3B-F), perhaps because control axonal mitochondria are already highly spherical. Interestingly, while the fission deficient mitochondria were longer in both axons and dendrites of *Drp1-mut* flies, their mitochondrial volume was either not different from controls in dendrites (Fig. S3G, H) or was significantly lower in axons (Fig. S3I, J); the latter was perhaps because of a lower diameter (Fig. 1G, S3E), most likely due to spatial constraints. The data overall show a highly significant and expected change in mitochondrial morphology in the fission-fusion mutant clock neurons and therefore indicate that this strategy can be used to assess the neuronal roles of mitochondrial fission and fusion.

### Mitochondrial fragmentation through loss of *Marf* but not *Opa1* leads to age-dependent loss of neuronal structure and function

We next used a membrane bound RFP to label neurons and their processes and assessed the structure and function of clock neurons with defective mitochondrial dynamics in young and old flies. Mitochondrial fission-deficient *Drp1-mut* clock neurons exhibited control-like neuronal morphology (Fig. 2A, S4A) and number of neurons (Fig. 2B). There was a similar lack of change in fusion-deficient *Opa1-mut* clock neurons and in young fusion-deficient *Marf-mut* clock neurons (Fig. 2A, S4A). However, some neurodegeneration was observed in older *Marf-mut* clock neurons accompanied by a small and significant reduction in neuron numbers (Fig. 2A, B).

**Fig. 2:**
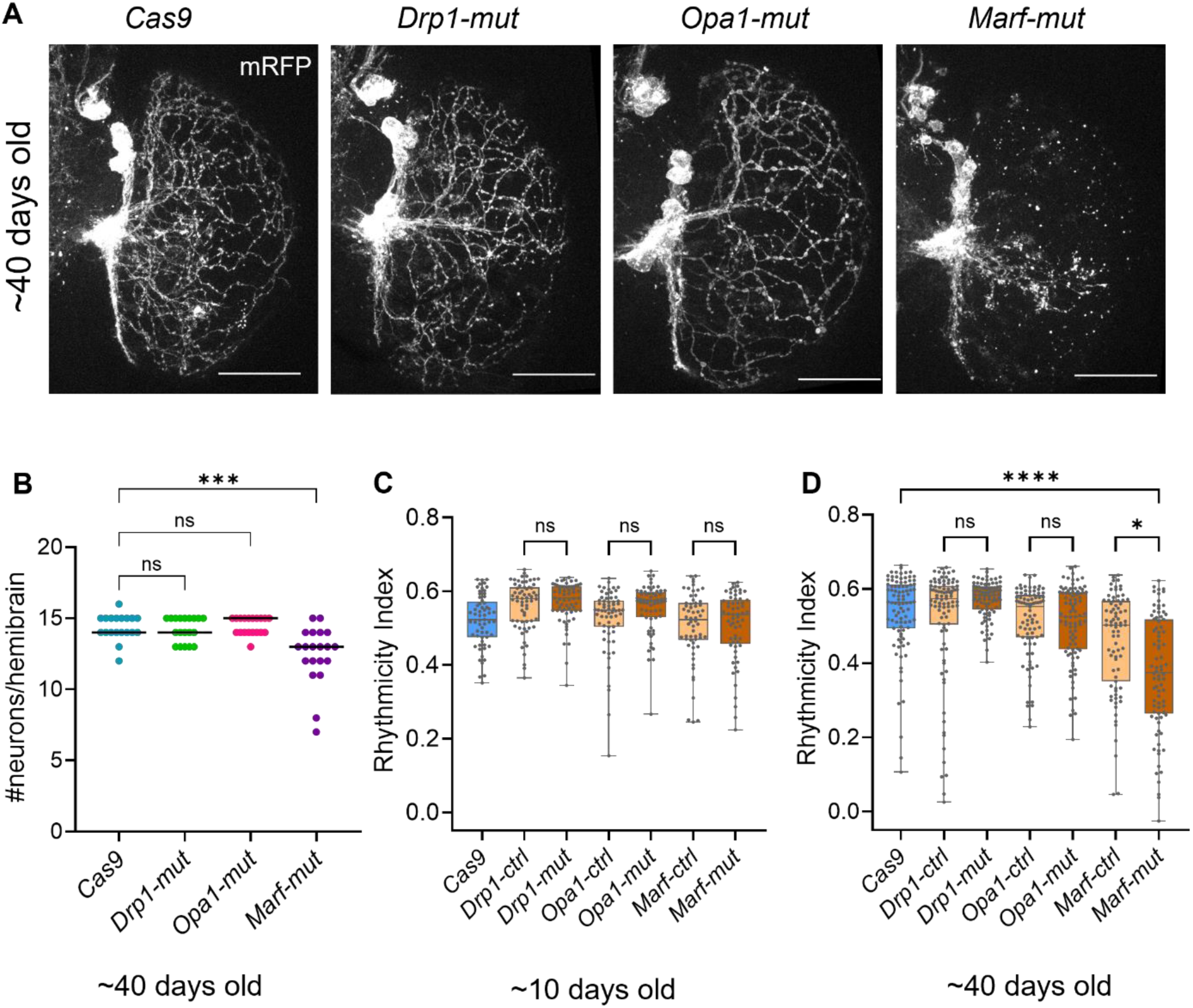
Marf-mutant clock neurons have age-dependent loss of structure and function. **A.** Representative images of ventral clock neurons and their projections from old flies labeled by *UAS-mRFP* expressed by *CLK856-Gal4.* Scale bars represent 50µm. Neurodegeneration can be seen only in *Marf-mut* old clock neurons (also see Fig. S4) **B.** Quantification of the number of ventral clock neurons (sLNvs, lLNvs and LNds) per hemibrain of 37-42 days old fly brains of indicated genotypes, n ≥ 19 hemibrains per genotype from two independent experiments. Kruskal-Wallis ANOVA with a post hoc Dunn’s test was used to test statistical significance between Cas9-control and the mutants, adjusted p-value >0.05 is denoted as not significant (n.s.); ***p<0.001. **C-D.** Rhythmicity Index, an indicator of circadian rhythm strength of flies with indicated mutations driven by *CLK856-Gal4*. Flies were loaded on to behavior experiments at ∼10 days old (C) or ∼40 days old (D), entrained in 12 hours light: 12 hours dark for at least three days before switching to constant darkness (DD) and Rhythmicity Index measured for DD days 2-7, n ≥ 60 flies per genotype from at least two independent experiments. Kruskal-Wallis ANOVA with a post hoc Dunn’s test was used to test statistical significance between mutants and Cas9-control as well as their respective gRNA controls, adjusted p-value >0.05 is denoted as not significant (n.s.); *p<0.05; ****p<0.0001.

Since both the molecular clock and neuronal firing in these neurons is required for circadian behavior (Schlichting et al., 2019), we used circadian rhythmicity as an initial assay of clock neuron function. Surprisingly, fission deficient *Drp1-mut* or fusion-deficient *Opa1-mut* clock neurons did not affect rhythmicity in young or old flies despite the dramatic changes in mitochondrial morphology (Fig. 2B-C). Both these perturbations did however lead to a small increase in circadian period, especially in older flies (Fig. S4B, C). In contrast, fusion-deficient *Marf-mut* clock neurons did show an age-dependent loss of neuronal function as indicated by an age-dependent decrease in rhythmicity (Fig. 2B-C). Together, these data indicate a bigger role for mitochondrial fusion mediated by *Marf* than by *Opa1* or than by mitochondrial fission in maintaining long-term neuronal structure and function.

### Transcriptomic analysis reveals an opposite effect of fission and fusion on mitochondrial transcripts

To further understand the cellular effects of perturbed mitochondrial dynamics in neurons including the different phenotypes of *Opa1-mut* and *Marf-mut* flies, we performed transcriptomic analysis of FACS-purified young and old clock neurons with and without the fission-fusion deficits (Fig. 3A). Loss of fission *Drp1-mut* neurons did not alter the transcriptome in young clock neurons and led to only a modest change in old neurons, with more genes upregulated than downregulated (Fig. 3B, C, D). Loss of fusion by *Marf-mut* also led to an age-dependent change in the transcriptome but with more genes downregulated than upregulated (Fig. 3B, E, F). Most surprisingly given its more modest phenotype, loss of fusion by *Opa1-mut* had the biggest number of transcriptomic changes in young as well as old clock neurons (Fig. 3B, H, I).

**Fig. 3:**
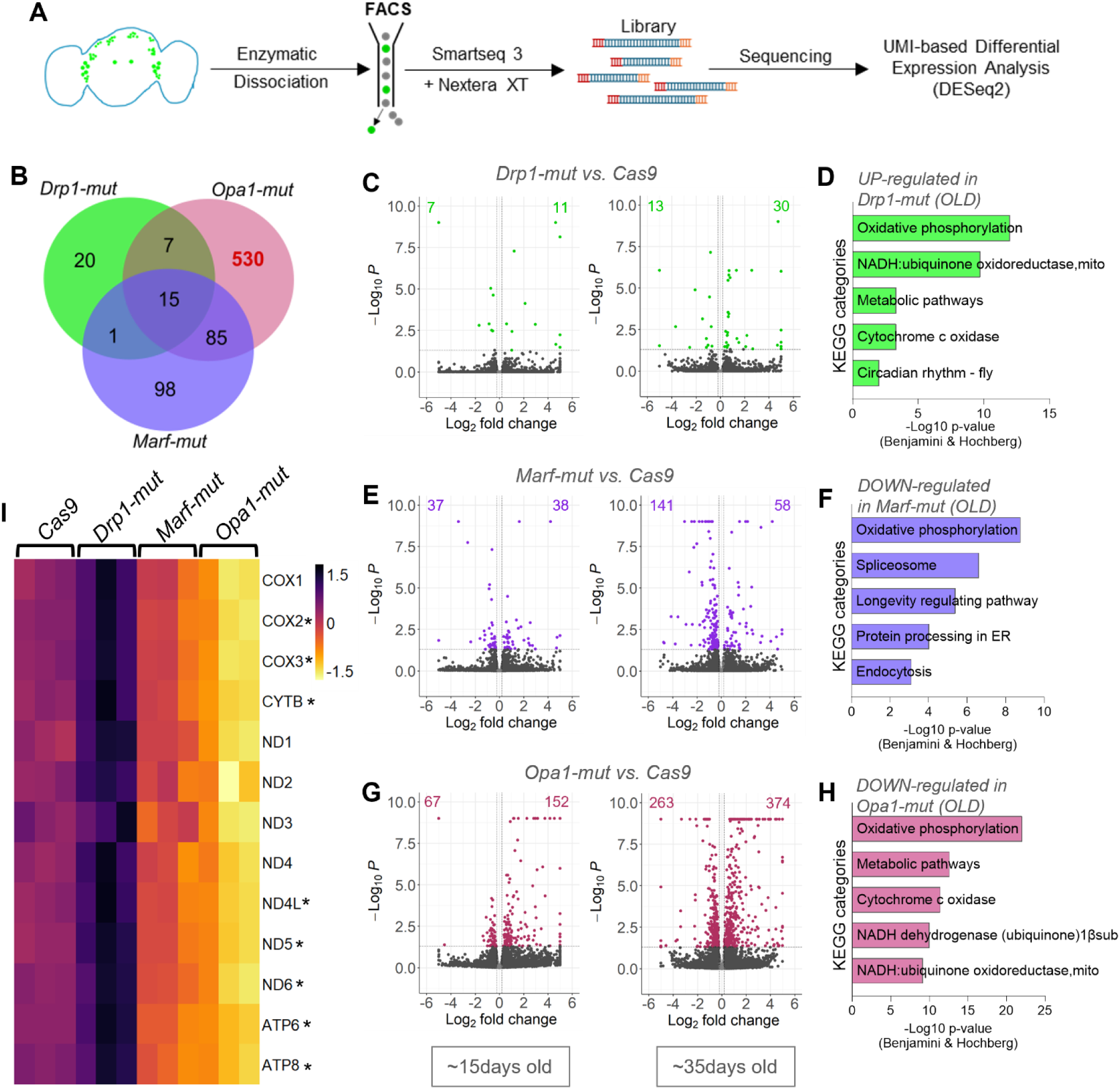
Loss of mitochondrial fusion leads to loss of mitochondrial transcripts in old clock neurons. **A.** Schematic representation of transcriptomic analysis from purified neurons. Clock neurons were labeled using *CLK856-Gal4>UAS-eGFP* apart from perturbations with Cas9 and gRNA. This strategy was applied for young (∼15 days old) and old (∼35 days old) flies and three replicates used per condition. **B**. Venn-diagram of all differentially expressed genes identified by DESeq2 (see methods) between the three perturbations in old clock neurons – *Opa1-mut* has the biggest change. **C,E,G.** Volcano plots of young (left) and old (right) mutant clock neurons compared to Cas9 control. Colored dots indicate significantly differentially expressed genes (adjusted p-value<0.05), number of genes up-regulated and down-regulated is indicated on each volcano plot. **D,F,H.** Significantly enriched categories by Gene Ontology classification of genes differentially regulated in old clock neurons by KEGG pathway. **I.** Heatmap showing expression levels of genes encoded by the mitochondrial genome in fission-fusion deficient clock neurons. All genes encoded by the mtDNA are down-regulated in *Opa1-mut* clock neurons, genes marked by an asterisk are significantly altered in all three conditions – upregulated in the fission mutant and downregulated in the fusion mutants.

Gene ontology analysis of differentially expressed genes identified oxidative phosphorylation-related genes to be most significantly upregulated in fission-deficient *Drp1-mut* clock neurons and downregulated in fusion-deficient *Marf* and *Opa1-mut* clock neurons (Fig. 3D, F, H). Mitochondria-associated, mtDNA-encoded transcripts were differentially regulated in an opposite manner by loss of fission and fusion (Fig. 3I, Fig. S5A). These transcriptional changes likely reflect a change in the mitochondrial DNA content as well as the ability of these neurons to produce ATP through oxidative phosphorylation (see Discussion).

### Chronic loss of Opa1 leads to a cancer-like transcriptomic response

Fusion-deficient *Opa1-mut* clock neurons that appear otherwise structurally and functionally healthy (Fig. 2) had the biggest transcriptomic changes, especially in old clock neurons in which, 263 genes were downregulated (Fig. 3B, G). They mainly contribute to oxidative phosphorylation, many more than those commonly downregulated with *Marf-mut* clock neurons (Fig. S5A, Fig. S5B). Most significantly, 374 genes were upregulated (Fig. 3G). They belong to several gene ontology categories, the most significant of which is related to glycolysis (Fig. 4A-C); this is reminiscent of a Warburg-like effect. In addition, genes related to protein translation initiation, elongation and aminoacyl-tRNA biosynthesis were significantly upregulated in *Opa1-mut* old clock neurons (Fig. 4A-B, S6A-B). The other two processes with significantly upregulated genes in old *Opa1-mut* clock neurons were related to ER-stress and oxidation-reduction (Fig. 4B, S7A-B). Interestingly, many of these genes and gene categories are upregulated in some or many cancers (Chen and Cubillos-Ruiz, 2021; de la Parra et al., 2018; Sangha and Kantidakis, 2022; Takahashi et al., 2021; Wang et al., 2016), indicating that a long-term loss of *Opa1* in neurons produces a cancer-like transcriptomic profile (See Discussion).

**Fig. 4:**
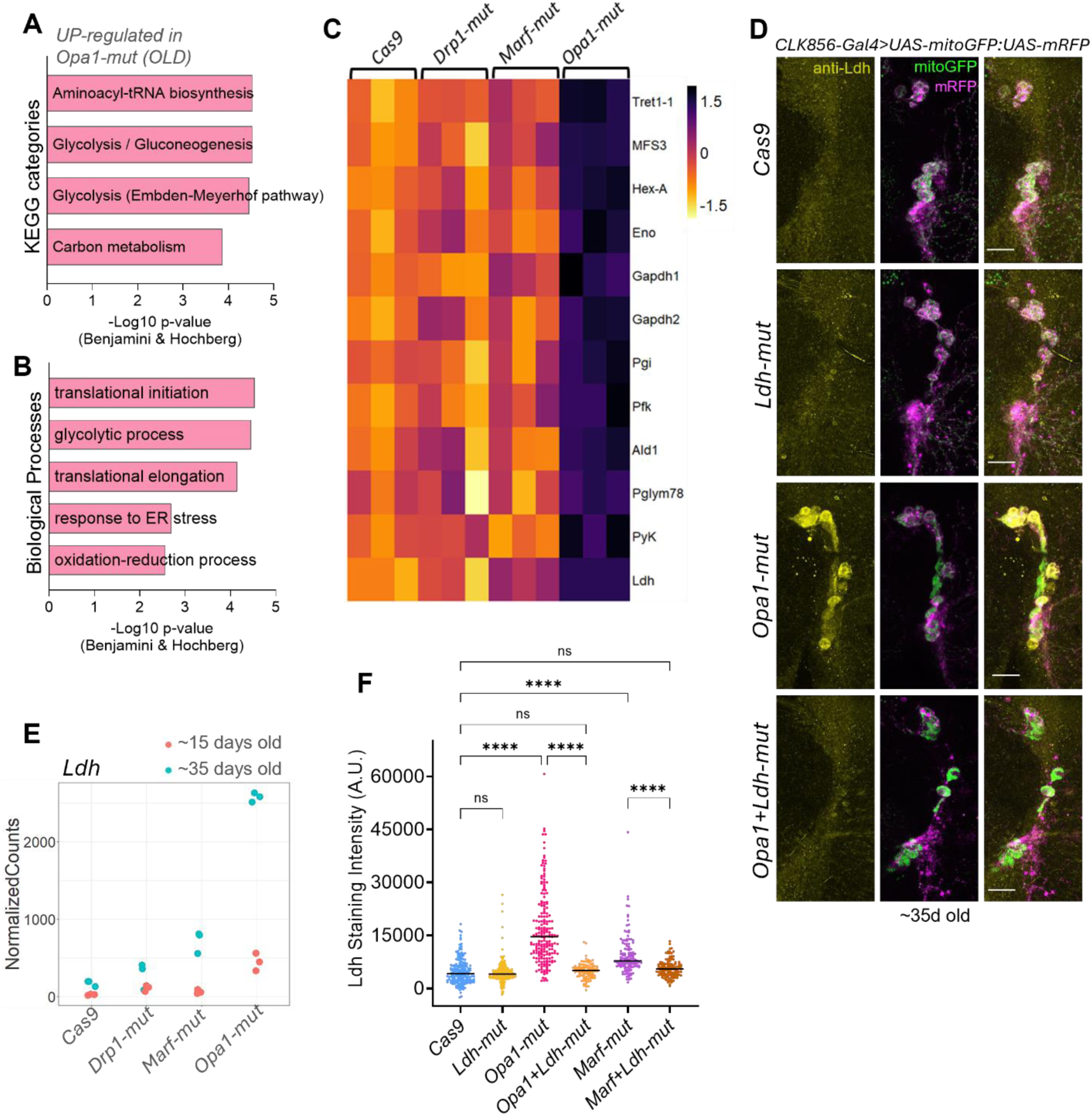
Loss of mitochondrial fusion through *Opa1* leads to Warburg-like effect in old clock neurons. **A-B.** Significantly enriched Gene Ontology classification of genes upregulated in old *Opa1-mut* clock neurons by KEGG pathway (A) and Biological Process (B). **C.** Heatmap showing expression levels of genes involved in the glycolysis pathway. Genes encoding for trehalose transporters – *Tret1-1* and *MFS3*, also included in the heatmap **D.** Representative images for LDH-staining in ventral clock neurons of indicated genotypes. Clock neurons were visualized using UAS-*mitoGFP* and UAS-*mRFP*, Scale bars represent 20 µm. **E.** Normalized counts for *Ldh* from the bulk sequencing dataset in young (pink) and old (cyan) clock neurons. **F**. Quantification of LDH-staining intensity from ventral clock neurons (sLNvs, lLNvs and LNds) in old flies, n ≥ 94 neurons from 12 or more hemibrains per genotype. Kruskal-Wallis ANOVA with a post hoc Dunn’s test was used to test statistical significance between groups, adjusted p-value >0.05 is denoted as not significant (n.s.); ****p<0.0001.

### Ldh-upregulation is neuroprotective in neurons deficient in mitochondrial fusion

One of the major hallmarks of cancer is the Warburg effect, the switching of cellular metabolism to aerobic glycolysis from mitochondrial oxidative respiration (Warburg, 1956). This is characterized by upregulation of genes involved in the glycolysis pathway and most importantly of Lactate dehydrogenase (*Ldh*) (Mishra and Banerjee, 2019). In old *Opa1-mut* clock neurons, there was significant upregulation of glycolysis-related genes (Fig. 4C), but the most dramatic change was in *Ldh* transcript levels. *Ldh* expression was upregulated ∼16 fold in old neurons (Fig. 4E). Similarly, there was *Ldh* upregulation in old *Marf-mut* clock neurons which are also fusion-deficient (∼4 fold), but to a lesser extent than in *Opa1-mut* neurons (Fig. 4E). The upregulation in old clock neurons was also apparent at the protein level by staining with an *Ldh* antibody (Fig. 4D, F).

We next investigated if this impressive transcriptomic change in *Ldh* is physiologically relevant and whether it is detrimental or compensatory. To this end, we employed a cell type-specific double guide strategy, i.e., we added an additional set of UAS-3X-guide RNA against *Ldh (Ldh-mut)* to the clock neuron-specific mitochondrial dynamic guides. Although *Ldh-mut* clock neurons did not alter rhythmic behavior, double mutants of *Opa1* and *Ldh* in clock neurons resulted in substantially reduced rhythmicity even in young flies (Fig. 5A). This became progressively worse with age (Fig. 5B), indicating a substantial age-dependent decrease in neuronal function in these double mutants. There was a similar but weaker interaction between *Ldh* and *Marf* (Fig. 5C), whereas there was no interaction with the fission gene *Drp1* (Fig. S8).

**Fig. 5:**
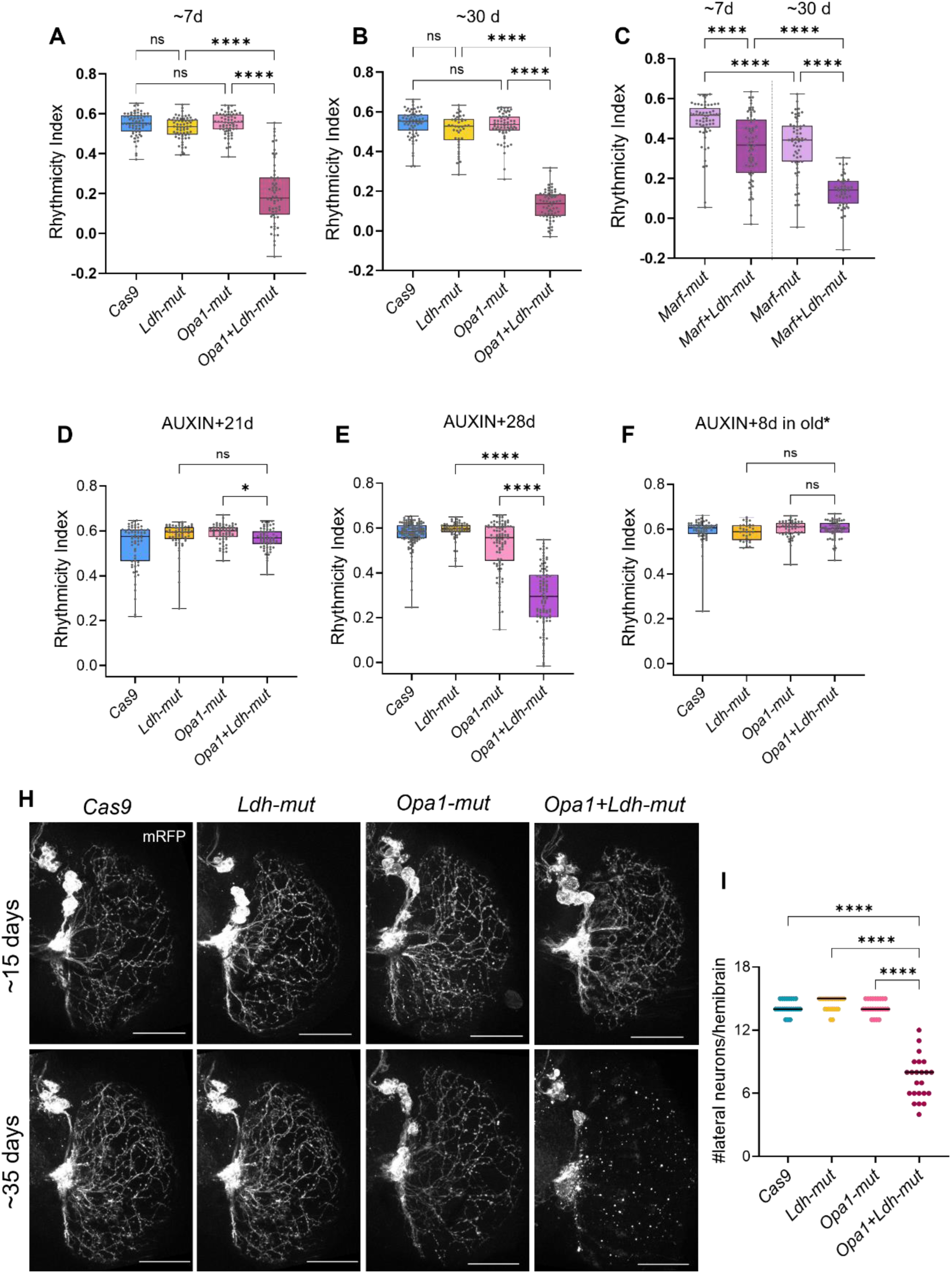
*Ldh* upregulation is protective in *Opa1*-deficient neurons. **A-C.** Rhythmicity of young (∼7 days old) and old (∼30 days old) with indicated mutations driven by *CLK856-Gal4* measured as described (Fig. 2 legend); n ≥ 41 flies per genotype from two independent experiments. Additional loss of *Ldh* in fusion deficient neurons leads to age-dependent loss of rhythmicity. Kruskal-Wallis ANOVA with a post hoc Dunn’s test, adjusted p-value >0.05 is denoted as not significant (n.s.); ****p<0.0001. **D-E.** Rhythmicity Index of flies containing AGES, an auxin inducible expression system combined with *CLK856-Gal4* that were fed auxin for 5 days and tested after the number of days indicated on the graphs, n ≥ 62 flies for each genotype from at least two independent experiments **F.** Rhythmicity Index for flies that were fed auxin for 5 days when old (*starting at ∼21 days and tested 8 days after), n ≥ 32 flies per genotype. Only in flies 28 days post auxin feeding, *Opa1+Ldh-mut* had significantly lower rhythmicity. Kruskal-Wallis ANOVA with a post hoc Dunn’s test, adjusted p-value >0.05 is denoted as not significant (n.s.); *p<0.05; ****p<0.0001 (See also Fig. S9) **H.** Representative images of ventral clock neurons and their projections labeled by *UAS-mRFP* expressed by *CLK856-Gal4* from young (∼15d) and old (∼35d) flies. Neurodegeneration can be seen in old *Opa1+Ldh-mut* clock neurons. Scale bars represent 50µm. **I.** Quantification of the number of ventral clock neurons (sLNvs, lLNvs and LNds) per hemibrain of old (∼35d) flies of indicated genotypes. *Opa1+Ldh-mut* had significantly lower number of neurons compared to either single mutant, Kruskal-Wallis ANOVA with a post hoc Dunn’s test, adjusted p-value ****p<0.0001.

Is the stronger phenotype of the *Opa1+Ldh* double mutant in older flies the result of the longer duration of the perturbation or the higher sensitivity of older neurons? To address this question, we used adult-specific expression system inducible by auxin feeding (AGES, (McClure et al., 2022)); we have previously shown that AGES functions well in clock neurons (Richhariya et al., 2023). Although mitochondrial morphology is altered in an expected manner at 11 days post auxin feeding in *Opa1+Ldh* double mutants (Fig. S9A-B), there is no effect on rhythmicity until 4 weeks (Fig. 5D, E, S9C-D). Moreover, feeding older flies auxin does not affect rhythmicity (Fig. 5F). In addition, *Opa1+Ldh* double mutant clock neurons without AGES undergo age-dependent neurodegeneration. Young double mutant clock neurons had neuronal numbers and process morphology similar to controls, but older ones suffered dramatic neurodegeneration of neuronal processes as well as a significant reduction in neuronal numbers (Fig. 5H, I). The data indicate that chronic loss of *Opa1+Ldh* expression impairs neuronal integrity and function.

### ATF4 regulates the up regulation of *Ldh* in response to loss of *Opa1* function

To further address the cellular and molecular response to disrupted mitochondrial fusion, we scanned for potential transcription factors that could orchestrate the observed upregulation. The transcriptomic data suggested 4 transcription factors that are upregulated in old *Opa1-mut* clock neurons and could potentially mediate this response (Fig. 6B, S10): *sima* (*Drosophila* ortholog of the hypoxia-inducible factor *HiF1α*) and *c-Myc* are known regulators of *Ldh* transcription in *Drosophila* and mammalian tumors (Firth et al., 1995; Shim et al., 1997; Wang et al., 2016; Wong et al., 2019); *kay* (*Drosophila* ortholog of the cFos gene) is a known regulator of circadian function (Ling et al., 2012); *crc* (*Drosophila* homolog of the mammalian ATF4 gene, henceforth referred to as ATF4) is involved in the integrated stress response (ISR) and is also upregulated in many tumors (Pakos-Zebrucka et al., 2016; Wortel et al., 2017). Moreover, ATF4 was identified as a master regulator of response to mitochondrial stress in mammals (Quirós et al., 2017) and a tumorigenic Warburg-effector in *Drosophila* epithelial cells (Sorge et al., 2020). We therefore hypothesized that loss of mitochondrial fusion via *Opa1* knockdown results in mitochondrial stress. It would upregulate *ATF4* expression, which in turn would upregulate other relevant genes like *Ldh*. We tested for *Ldh* upregulation using a double mutant for *Opa1* and *ATF4*. Indeed, this genetic combination failed to upregulate *Ldh* expression to the same extent as single *Opa1-mut* (Fig. 6A, C) and showed age-dependent loss of rhythmicity (Fig. 6D, E). Similar to the *Opa1+Ldh* double mutants (Fig. 5H), *Opa1+ ATF4* double mutants also undergo age-dependent neurodegeneration (Fig. 6F). We conclude that ATF4 mediates the neuroprotective upregulation of *Ldh* in fusion-deficient clock neurons.

**Fig. 6:**
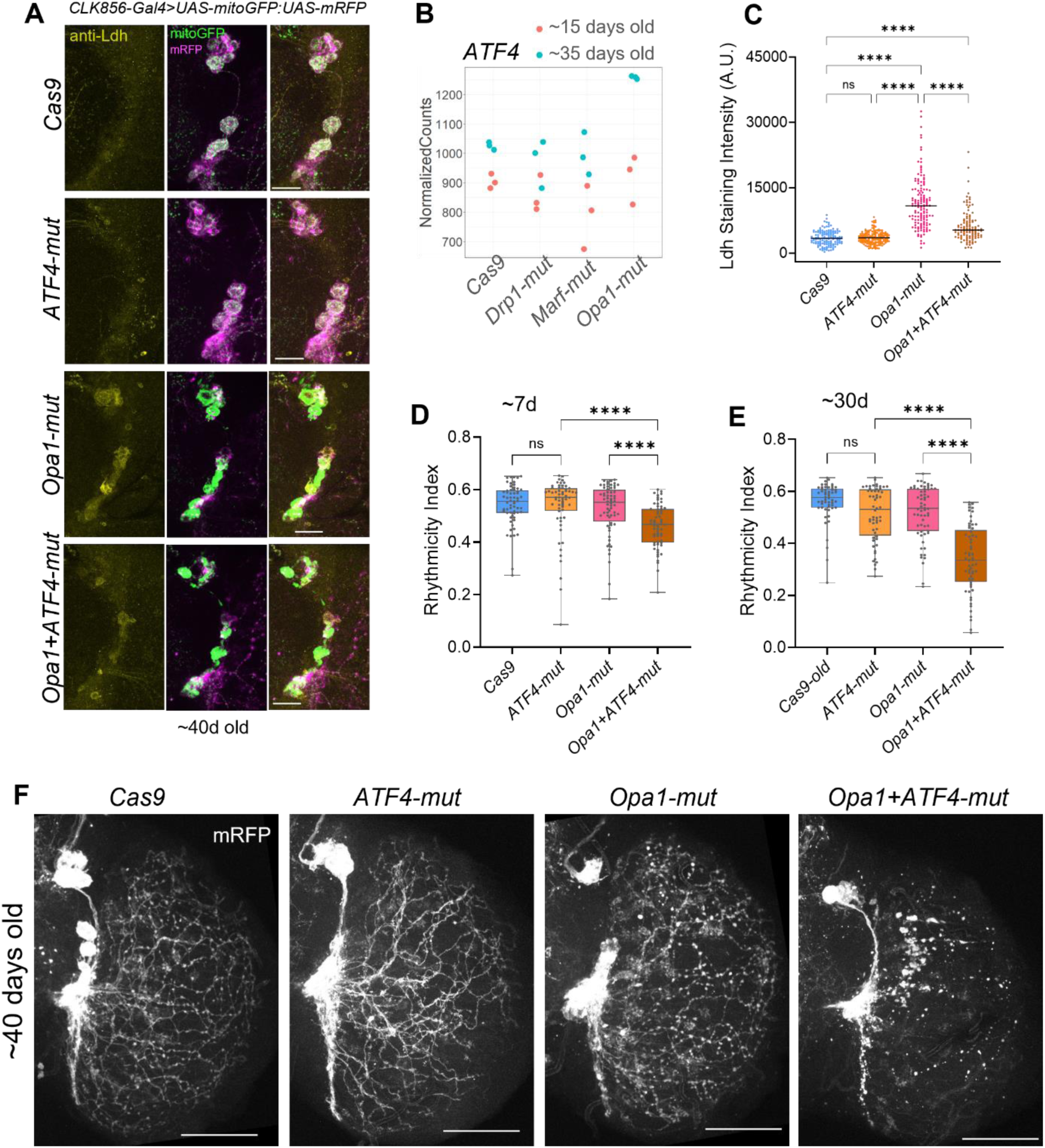
*Ldh* upregulation is mediated by ATF4. **A.** Representative images for LDH-staining in ventral clock neurons of the indicated genotypes. Clock neurons are labeled by UAS-*mitoGFP* and UAS-*mRFP* Scale bars represent 20 µm. **B.** Normalized counts for *ATF4* from the bulk sequencing dataset in young (pink) and old (cyan) clock neurons. **C.** Quantification of LDH-staining intensity from ventral clock neurons (sLNvs, lLNvs and LNds) in old flies, n ≥ 110 neurons from 11 or more hemibrains per genotype. *Opa1+ATF4*-mutant clock neurons could not upregulate LDH to the same extent as *Opa1-mut*, Kruskal-Wallis ANOVA with a post hoc Dunn’s test, adjusted p-value >0.05 is denoted as not significant (n.s.); ****p<0.0001 **D-E.** Rhythmicity Index of young (∼7d old) and old (∼30d old) flies with indicated mutations driven by *CLK856-Gal4* measured as described (Fig. 2 legend); n ≥ 54 flies per genotype from two independent experiments. Loss of rhythmicity is seen at both ages in *Opa1+ATF-double mutant* clock neurons, Kruskal-Wallis ANOVA with a post hoc Dunn’s test, adjusted p-value >0.05 is denoted as not significant (n.s.); ****p<0.0001 **F.** Representative images of ventral clock neurons and their projections labeled by *UAS-mRFP* expressed by *CLK856-Gal4* of ∼40day old brains. Neurodegeneration is evident in *Opa1+ATF4-mut* clock neurons. Scale bars represent 50µm.

## Discussion

To study the contribution of mitochondrial fission and fusion to neuronal function, we used a cell type-specific CRISPR approach to knockout key fission-fusion genes and disrupt normal mitochondrial dynamics of *Drosophila* clock neurons. Simple morphological comparisons with wild-type mitochondria indicate that these knockouts appear to completely eliminate mitochondrial fission or fusion, within clock neuron axons and dendrites as well as cell bodies (Fig.1, S2, S3). With these hyperfused and fragmented mitochondria, we thought that the morphological assays might indicate some circadian effect on wild-type mitochondrial morphology, but there was no dramatic change in the mix of spherical and tubular mitochondria at different times of day (data not shown).

Surprisingly, the chronic absence of fission or fusion had no detectable effect on rhythmicity or on circadian period in many genotypes and circumstances (Fig. 2), suggesting little or no effect on neuronal function. Some of these results were verified by GCaMP assays, i.e., there was little or no effect of these chronic knockouts on the calcium response of clock neurons to *in vitro* carbachol stimulation (data not shown). One interpretation is that the knockouts have no effect on neuronal activity. Another is that behavioral assays do not provide a sufficiently sensitive readout of activity. For example, the knockouts might lower neuronal activity but it is still above a threshold that provides wild-type-like behavior. There were however a few genotypes and circumstances with clear behavioral phenotypes, which indicate mitochondrial effects on neuronal function especially in older flies. Moreover, the differences between fission and fusion knockouts were interesting.

### Mitochondrial fusion is more important than fission in adult neurons

We were surprised by the modest contribution of blocking fission to neuronal health and function despite its dramatic effect on mitochondrial morphology (Figs. 1&2). The larger effect of blocking fusion suggests an overall more important role, especially in aging *Drosophila* neurons. A similar more important role for fusion was identified in *C. elegans* neurons and muscles (Byrne et al., 2019). Several diseased conditions manifest fragmented mitochondria (Lutz et al., 2009; Wang et al., 2009, 2008; Wang and Davis, 2021), indicating their association with dysfunction. Perhaps this mostly reflects a more important contribution of fused than fragmented mitochondria to normal function.

Nonetheless, mitochondrial fission is known to have a role in aging. For example, increasing mitochondrial fission ubiquitously in midlife promotes healthy aging in flies (Rana et al., 2017), whereas hyperfused mitochondria promote senescence in mice (Yu et al., 2020). Additionally, mitophagy-associated fission is important for mitochondrial function in aging worms and mice (D’Amico et al., 2019) . However, none of these perturbations were neuron-specific. It is therefore likely that many post-mitotic neurons with very limited mitochondrial fission like the clock neurons shown here are healthy and functional, perhaps in part because mitophagy can be independent of mitochondrial fission (Anzell et al., 2021; Yamashita et al., 2016). As a caveat, we note that our results here are limited to clock neurons under normal conditions, meaning that mitochondrial fission might be more important in other neurons or under other conditions like stress. In any case, this study suggests that enhanced mitochondrial fragmentation can have diverse causes and consequences and should therefore be studied more carefully.

### Gene expression responses to clamping mitochondrial fission-fusion

Oxidative phosphorylation-related genes are the most significantly upregulated in *Drp1-mut* neurons and downregulated in fusion-deficient *Marf-mut* and *Opa1-mut* neurons (Fig. 3D, F, H). The simplest interpretation is that these changes reflect the fraction of fused mitochondria, which probably engage in more potent oxidative phosphorylation than fragmented mitochondria due to their higher mRNA levels. Indeed, mitochondria-encoded transcripts were also among those differentially regulated in an opposite manner by loss of fission and fusion, up in *Drp1-mut* neurons and down in *Marf-mut* and *Opa1-mut* neurons (Fig. 3I). These changes might predominantly reflect changes in mitochondrial DNA content, less in fragmented than fused mitochondria (Chen et al., 2007). The larger number of changes in oxidative phosphorylation-related genes in the *Opa1-mut* neurons might also reflect differences in mitochondrial DNA content but more indirectly than the mitochondria-encoded transcripts.

Opa1 and Marf both promote fusion but they do so in different ways: Opa1 mediates fusion of the inner mitochondrial membrane, whereas Marf promotes fusion of the outer membrane. The bigger gene expression change from inhibiting Opa1 function might reflect its more intimate association than Marf with the electron transport chain via the inner mitochondrial membrane. Indeed, Opa1 plays a role in maintaining cristae structure in mammalian cells in addition to its role in mitochondrial fusion (Patten et al., 2014). It is therefore likely that loss of Opa1 induces additional mitochondrial damage beyond the loss of fusion, which then triggers a robust Warburg effect. This gene expression difference might also explain the quite big difference in behavioral phenotype between the two fusion factor knockouts: the more modest upregulation of the Warburg factors and especially LDH by Marf inhibition (Fig. 4E, F) might leave these neurons more functionally impaired than by Opa1 inhibition.

### Comparisons with non-neuronal cells and mammalian systems

Different modes of mitochondrial fragmentation in *Drosophila* cardiomyocytes similarly resulted in different cellular effects: Drp1 overexpression had no effect; Opa1 inhibition was associated with ROS; Marf was associated with ER-stress (Bhandari et al., 2015). Similarly, loss of the mammalian ortholog of Marf in POMC neurons led to loss of function mediated by ER stress (Schneeberger et al., 2013). These Marf effects probably reflect the association between the ER and mitochondria via its outer mitochondrial membrane. Some toxins can also fragment mitochondria (Passmore et al., 2017). Note that environmental factors such as high fat diet can induce mitochondrial fragmentation by downregulating genes involved in mitochondrial fusion (Zheng et al., 2023).

### Neurodegeneration and the role of Ldh in preventing neurodegeneration

Although some studies have explored the interaction of mitochondrial fragmentation due to Drp1 hyperactivation with neurodegeneration (Cho et al., 2009; Petrozziello et al., 2022), these effects could be due to a reduction in fusion, and genes associated with mitochondrial fusion are also downregulated in some neurodegenerative diseases (Shirendeb et al., 2011; Wang et al., 2008). Indeed, the loss of fusion is a major link to neurodegeneration at least as shown here in the absence of *Opa1* and *Ldh*. In this context, the double mutant phenotype indicates that *Ldh* contributes to a remarkable level of compensation (Fig. 5A, B), likely due to the dramatic upregulation in *Ldh* levels (Fig. 4D). This is because *Opa1*-single mutant clock neurons function almost like controls with only a minor effect on circadian period even in old clock neurons (Fig. 2, S4). Interestingly, *Ldh* is also upregulated in different *Drosophila* neurons under other conditions of mitochondrial stress (Granat et al., 2023; Hunt et al., 2019).

There are a few possible mechanisms of *Ldh* rescue of fusion-deficient neurons, the most obvious one being via ATP levels. Ldh can generate ATP via glycolysis in the absence of mitochondrial function albeit at a much lower efficiency than oxidative phosphorylation (Valvona et al., 2016). Fusion mutants in liver, heart and muscle also lose mitochondrial DNA (Chen et al., 2010, 2007, 2012; Lee et al., 2023), which likely leads to lower mitochondrial function and ATP production as is likely the case in *Opa1-mut* and *Marf-mut* clock neurons (Fig. 3K).

In addition to generating ATP, *Ldh* also regenerates NAD from NADH, which allows for sustaining glycolysis. Interestingly, expression of the yeast NADH dehydrogenase Ndi1 can compensate for disrupted mitochondrial Complex I in different organisms (Granat et al., 2023; Han et al., 2023; Meisel et al., 2024). Although Ndi1 does not pump protons, these effects could still be caused by a least partial rescue of oxidative phosphorylation (Jiménez-Gómez et al., 2023). Lastly, fragmented mitochondria are associated with elevated ROS (Hung et al., 2018; Yu et al., 2006), due perhaps to less efficient oxidative phosphorylation. If *Ldh* upregulation in fusion mutants channels energy production mostly through aerobic glycolysis, this would reduce mitochondrial oxidative phosphorylation and as a consequence reduce reactive oxygen species (ROS) mediated oxidative damage and neurodegeneration. In summary, *Ldh* upregulation might protect Opa1-deficient neurons from either low ATP, low NAD or high ROS, or quite likely a combination of the three, which then leads to neurodegeneration.

These deficiencies might be greater in old neurons, or old neurons more vulnerable, thereby explaining why neurodegeneration is more prominent in old than young neurons. However, chronic mitochondrial fragmentation is required to cause dysfunction; acute fragmentation even in old neurons had no effect (Fig. 5D-F). This indicates the late-onset of neurodegeneration is likely a long-term effect from these deficiencies rather than any vulnerability of older neurons. More detailed timing experiments should clarify how rapidly neurodegeneration occurs. We also note that mammalian neurons have much longer lifespans than fly neurons, but the same approach we have taken here, acute vs chronic fragmentation, should be straightforward in mammalian systems.

### Relationship between neurodegeneration and cancer

The transcriptomic response seen in *Opa1*-mutant old clock neurons is strikingly similar to that observed in many cancers (Figs. 4, S6, 7). Although the initial hypothesis was that mitochondrial dysfunction was the primary cause of the Warburg effect, many cancers with a Warburg effect have normal mitochondrial function, and oncogenic triggers are now considered the primary reason for the shift to aerobic glycolysis (Vaupel and Multhoff, 2021). Fragmented mitochondrial morphology is positively correlated with proliferation in cancer cells (Chen and Chan, 2017). These data point to the potential similarity of molecular responses in tumors and neurodegenerative cells. However, the effects on disease prognosis appear to be opposite. In most cancers, upregulation of glycolysis is tumorigenic, and elevated *Ldh* levels are associated with a poor prognosis (Comandatore et al., 2022; Wulaningsih et al., 2015). In contrast, we show here that *Ldh* and likely elevated glycolysis are highly neuroprotective. Interestingly, incidences of cancer and neurodegeneration are anti-correlated in the general population for reasons that are not well understood (Houck et al., 2018; Plun-Favreau et al., 2010). The opposite effects of *Ldh-*upregulation on disease prognosis of cancer and neurodegeneration is a possible mechanism.

### A model to more generally explore the genetics of neurodegeneration

Many neurodegenerative diseases are described based on clinical symptoms or post-mortem brain samples. This is partly because the genetic basis of neurodegeneration is complex. For several neurodegenerative diseases such as Parkinsons, Alzheimers and ALS, monogenic or mendelian causes explain only 5-20% of the cases; the others are idiopathic and sporadic (Bandres-Ciga et al., 2020; Ghasemi and Brown, 2018; Hoogmartens et al., 2021). Sporadic cases have several known genetic risk factors, but most have low penetrance, and there are also environmental influences. Mitochondrial dysfunction is among the targets of these genetic and environmental factors (Tran et al., 2020).

Mitochondrial dynamics and function are highly conserved between *Drosophila* and mammalian cells. Indeed, the two major interactors of mitochondrial fusion we identified in context of neurodegeneration, *Ldh* and *ATF4* (Fig 5, 6), are likely relevant to mammalian neurodegeneration. Indeed, lactate levels are reported as elevated in multiple human studies – although this finding has not always been replicated (Wang et al., 2024). As there are likely many other interactions that contribute to neurodegeneration or neuroprotection, the genetic model used in this study should facilitate enhancer-suppressor genetics and identify additional conserved factors that bridge mitochondrial function to aging and neurodegeneration.

## Materials and Methods

### Fly rearing and stocks

Flies were raised on standard cornmeal medium supplemented with yeast at 25°C under 12h light: 12h dark conditions. For experiments using aged-flies, ∼0-4 days old male flies were collected separated into vials containing 30-40 flies. Aging flies were transferred to fresh food every 3-4 days until ready for experiments. All fly lines used are listed in Table S1.

For auxin feeding in adult-specific experiments, adult male flies of the correct age were transferred to Instant *Drosophila* Medium (Formula 424®, Carolina) reconstituted in 10mM Auxin (Glentham Life Sciences #GK2088) in 1X PBS. Auxin feeding was done for 4-5 days with food change at 2-3 days.

### Generation of gRNA lines

Lines expressing 3X-gRNA under UAS-control were generated using previously described protocols (Port and Bullock, 2016; Schlichting et al., 2019). CRISPR Optimal Target Finder tool (Gratz et al., 2014) was used to identify gRNA specific to target genes. These sequences were incorporated in primers listed in Table S2 and used to amplify fragments from the pCFD6 vector (Addgene #73915). The final plasmid was assembled using NEBuilder^®^ HiFi DNA Assembly Cloning Kit (NEB), combining the two PCR fragments and pCFD6 vector linearized with BbsI-HF (NEB). Plasmid sequences were verified by Sanger sequencing. Plasmids were injected into embryos by Rainbow Transgenic Flies Inc (Camarillo, CA, USA). The *UAS-3x-gRNA* lines against *Drp1, Opa1* and *Marf* were inserted into the attP1 site on the second chromosome whereas the UAS-3x-gRNA line against *Ldh* was inserted into the attP2 site on the third chromosome. Positive transformants were screened for by red eye color based on the mini-white marker.

### Longevity assay

For each genotype, 0-4 days old male flies were collected and divided into ∼20 flies per vial. A total of at least 68 flies were used for each genotype. Flies were transferred every 3-4 days to fresh food while counting dead flies. Survival curves were plotted using GraphPad Prism.

### Circadian Behavior Assay

Circadian behavior was assayed as described previously (Richhariya et al., 2023). Briefly, male flies of the desired age were loaded into *Drosophila* Activity Monitor (DAM) tubes containing sucrose food (4% sucrose and 2% agar). Light boxes with programmable LED light intensities were used to control light conditions and were kept inside a temperature-controlled incubator set to 25°C. Flies were entrained to at least 3 days of a 12 hours light: 12 hours dark cycle before switching to constant darkness for at least 7 days. Rhythmicity index, used as a measure of the strength of the circadian rhythm was calculated for constant darkness (DD) days 2-7 using Sleep and Circadian Analysis MATLAB Program (SCAMP) developed by Christopher G. Vecsey (Donelson et al., 2012). Period values for only rhythmic flies (RI>0.3) were used.

### Immunohistochemistry

Immunostaining for *Drosophila* brains was performed as described previously (Richhariya et al., 2023). Briefly, flies were fixed in 4% PFA (Fisher Scientific #50-980-487) then brains were dissected and blocked in blocking buffer (10% Normal Goat Serum (NGS, Jackson Labs #005-000-121) in 0.5% PBST). The following primary antibodies in blocking buffer were used: chicken anti-GFP (1:2000, Abcam #ab13970), rat anti-RFP (1:1000, Proteintech #5f8), rabbit anti-LDH (1:500, Boster Bio #DZ41222). Primary antibody incubations were done for 15-18h at 4°C for GFP and RFP or ∼48h at 4ᴼC if LDH was used. Secondary antibodies were used at 1:500 dilution in blocking buffer and were incubated either for 3 hours at RT or 4°C overnight. All samples compared to each other were processed together. Brains were mounted in Vectashield mounting medium (Vector Laboratories #H-1000-10). Brains used for LDH quantification were mounted in Vectashield PLUS (Vector Laboratories #H-1900).

### Image acquisition and analysis

All images were acquired on Leica Stellaris 8 confocal microscope equipped with a white light laser. Imaging settings were constant across different samples in a set. Image processing and analysis were performed using Fiji (Schindelin et al., 2012).

For mitochondrial morphology analyses, images were acquired using a 63X oil objective with NA of 1.4 with additional optical zoom. Deconvolution was performed on acquired images using LIGHTNING on the LAS X software. Mitochondrial parameters were quantified using the Mitochondrial Analyzer plugin (Chaudhry et al., 2020) with Fiji. Images were analyzed using the batch 3D mode and parameters quantified on a per mito basis. For mean branch length, only non-zero values are reported.

For visualizing the projections of ventral clock neurons, images were acquired of the optic lobe region with the 20X air objective with an NA of 0.75 with additional optical zoom. Z-stack projections with maximum intensity are shown as representative images. Images were adjusted for brightness/contrast, with the same level of correction applied to all the samples in an experiment.

For quantification of LDH levels, images were acquired similarly as described above for visualizing projections. Images for both mRFP and LDH were acquired using sequential scans. For quantification, the Time Series Analyzer v3 Plugin with Fiji was used. Z-planes with cells were identified using the RFP channel and then LDH-levels were measured from the same pixels using the LDH-channel. Background was subtracted by measuring randomly selected areas outside of the cells.

### FACS-sorting of *Drosophila* neurons

Clock neurons labelled by CLK856-Gal4 and UAS-eGFP were purified using Fluorescence-activated cell sorting (FACS) using methods previously described (Schlichting et al., 2022). Briefly, male flies either 12-16 days old (∼15days old) or 34-38 days old (∼35 days old) at ZT14 were anesthetized on ice in the dark followed by dissections in small batches in ice-cold Schneider’s *Drosophila* medium (SM, Gibco #21720001). About 80-100 brains were dissected for each genotype and incubated in an enzyme solution containing 0.75ug/ul Collagenase (Sigma-Aldrich, #C1639) and 0.04 μg/ul Dispase (Sigma-Aldrich, #D4818) in SM for 40 minutes at RT. After washing and dissociation by trituration, a single cell suspension was filtered through a 40µm filter and the volume was adjusted to ∼2ml. Hoechst stain (Invitrogen, #R37605) was used to identify live cells. GFP-positive cells of interest were collected with gates that were set using a sample without GFP. ∼1000 GFP-positive clock neurons were collected per sample in 100µl lysis buffer (Dynabeads mRNA direct kit, Invitrogen #61011) and frozen immediately at -80°C. Three sets of samples were collected per genotype and age as replicates.

### Generation of sequencing libraries

PolyA mRNA was isolated using the Dynabeads mRNA direct kit (Invitrogen #61011) and eluted into 9ul Milli-Q water. 4.5ul of this polyA RNA was then used to make full-length cDNA with unique molecular identifiers (UMIs) using the Smartseq3 protocol. The original Smartseq3 single cell protocol (Hagemann-Jensen et al., 2020a, 2020b) was adapted for bulk neurons by using 5X volumes at each step combined with 15 PCR cycles. Amplified cDNA was purified using 0.6X volume of AMPure XP beads (Beckman Coulter #A63881) and checked for integrity and concentration using the High Sensitivity D5000 ScreenTape (Agilent # 5067-5592) on a TapeStation.

∼750pg of cDNA was then used as input into the Nextera XT DNA Library Preparation Kit (Illumina # FC-131-1096) with some modifications to prevent smaller fragment sizes. 0.8µl Amplicon Tagment Mix (ATM) was used with 9.2µl of cDNA+ 2X Tagmentation buffer + Milli-Q water. Tagmentation was performed at 55°C for 10 mins, quenched immediately by addition of 2.5µl buffer NT. On ice, 2.5ul each of the i5 and i7 indexes along with 7.5ul of NPM were added to make the final volume 25ul. 9 PCR cycles were used. Final libraries were purified using 0.7X volume of AMPure XP beads (Beckman Coulter #A63881) and quantified using a High Sensitivity D5000 ScreenTape (Agilent # 5067-5592) on a TapeStation.

These libraries were then sequenced on Illumina NovaSeq6000 at Novogene Corporation Inc. (Sacramento, CA, USA) generating 20-38million total raw paired end reads (2*150bp) per sample.

### Sequencing data analysis

The reads were mapped to the *Drosophila* genome (dm6) using the zUMIs pipeline (Parekh et al., 2018). All samples sequenced were processed simultaneously. UMI counts mapping to exons of each gene were extracted to be used for differential expression analysis.

Differential expression analysis was performed using DESeq2 (Love et al., 2014) using RStudio. Young and old samples were analyzed separately. Differentially expressed genes were identified by comparing the UMI counts of each of the mutants to the Cas9 controls of the same age. An adjusted p-value cutoff of <0.05 was used to define differentially expressed genes. No fold change cutoff was used. Gene Ontology enrichment analysis was performed using PANGEA (Hu et al., 2023) using genes expressed in clock neurons as a background set. Volcano plots were generated from the DESeq2 output using the package Enhanced Volcano (Blighe K, Rana S, Lewis M, 2023). DESeq2 output normalized counts were transformed using normTransform and then used to create heatmaps of selected genes with pheatmap (Kolde R, 2019). Heatmaps were scaled for rows and not clustered for rows or columns.

### Data Representation and Statistical Analysis

Behavior data are presented as box plots showing all data points, and whiskers extending from min to max. For quantification of mitochondrial properties as well as LDH staining intensity, all data points are plotted with median as a line. All statistical analysis other than sequencing data was performed using GraphPad Prism software version 10. For all of these experiments, samples were tested for normality, and if the distribution of any sample was not normal, the non-parametric Kruskal Wallis ANOVA was used with a post-hoc Dunn’s test. See figure legends for details.

## Acknowledgements

We thank Michelle Lin for assistance with maintaining fly stocks. We thank all members of the Rosbash lab for useful discussions. We thank Dr. Joshua Meisel for helpful comments on the manuscript. Fly stocks obtained from Bloomington *Drosophila* Stock Center (NIH P40OD018537) and Vienna Drosophila Resource Center (VDRC) were used in this study. This work was supported by the Howard Hughes Medical Institute (HHMI).

## Supplementary Figures

**Fig. S1:**
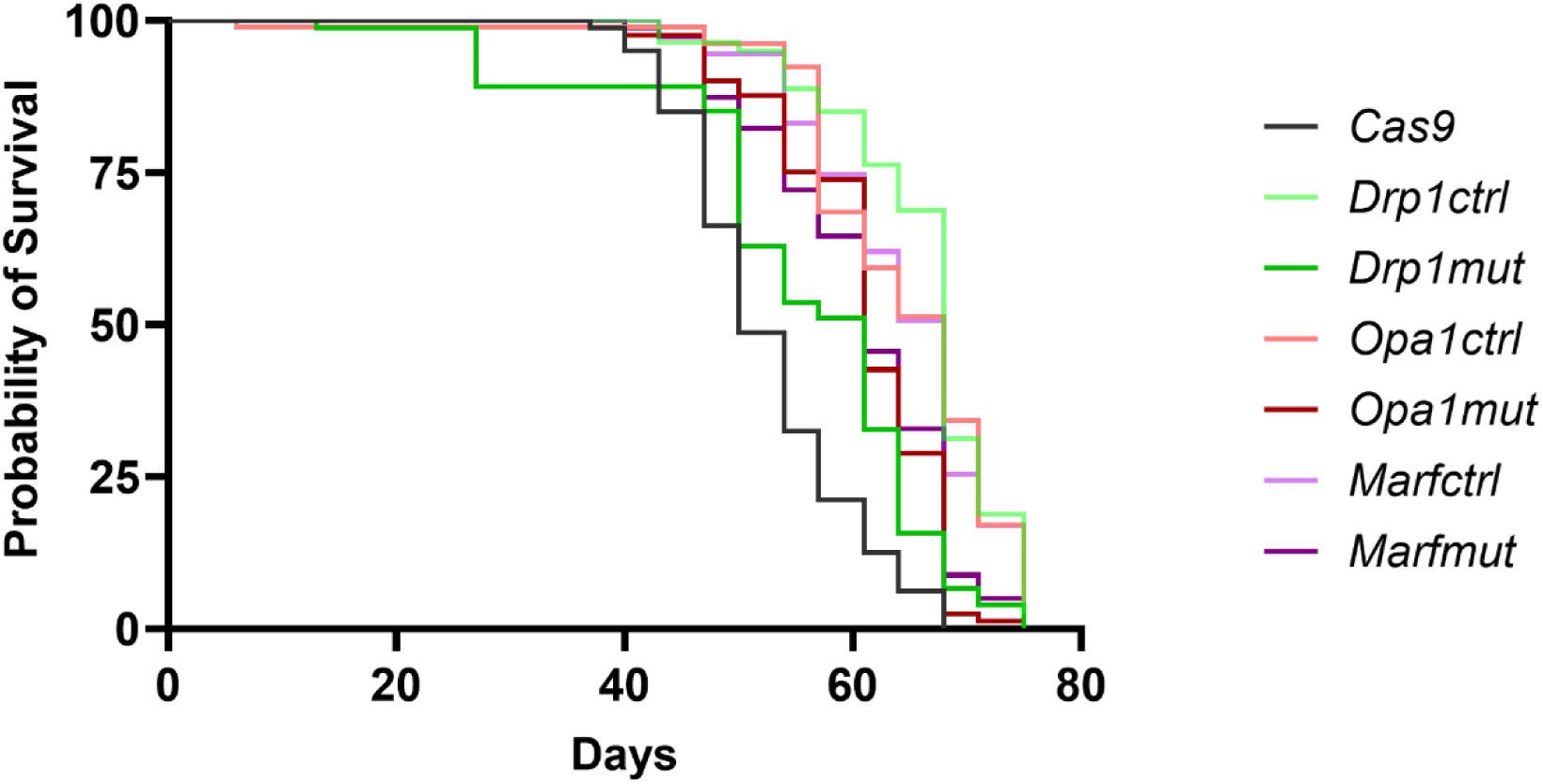
Loss of mitochondrial fission or fusion in clock neurons does not affect lifespan. None of the mutants survived for significantly lesser time than both Cas9 and the corresponding gRNA control (Log-rank Mantel-Cox test).

**Fig. S2:**
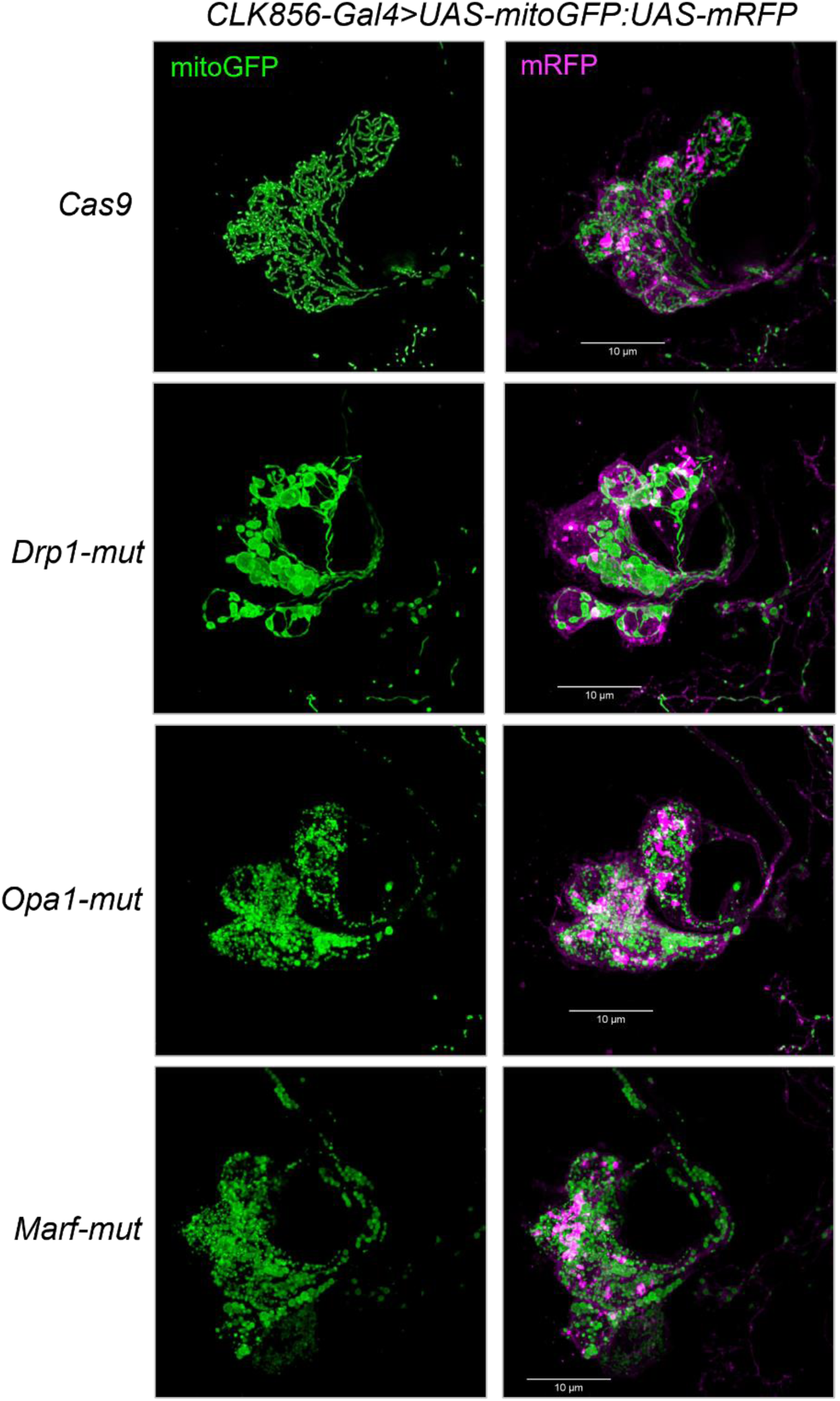
Mitochondrial morphology in the LNd clock neurons. Mitochondrial morphology in the LNd neuron cell bodies is altered as expected (Fig. 1B) using cell type-specific CRISPR with *CLK856-Gal4*. Scale bars represent 10µm.

**Fig. S3:**
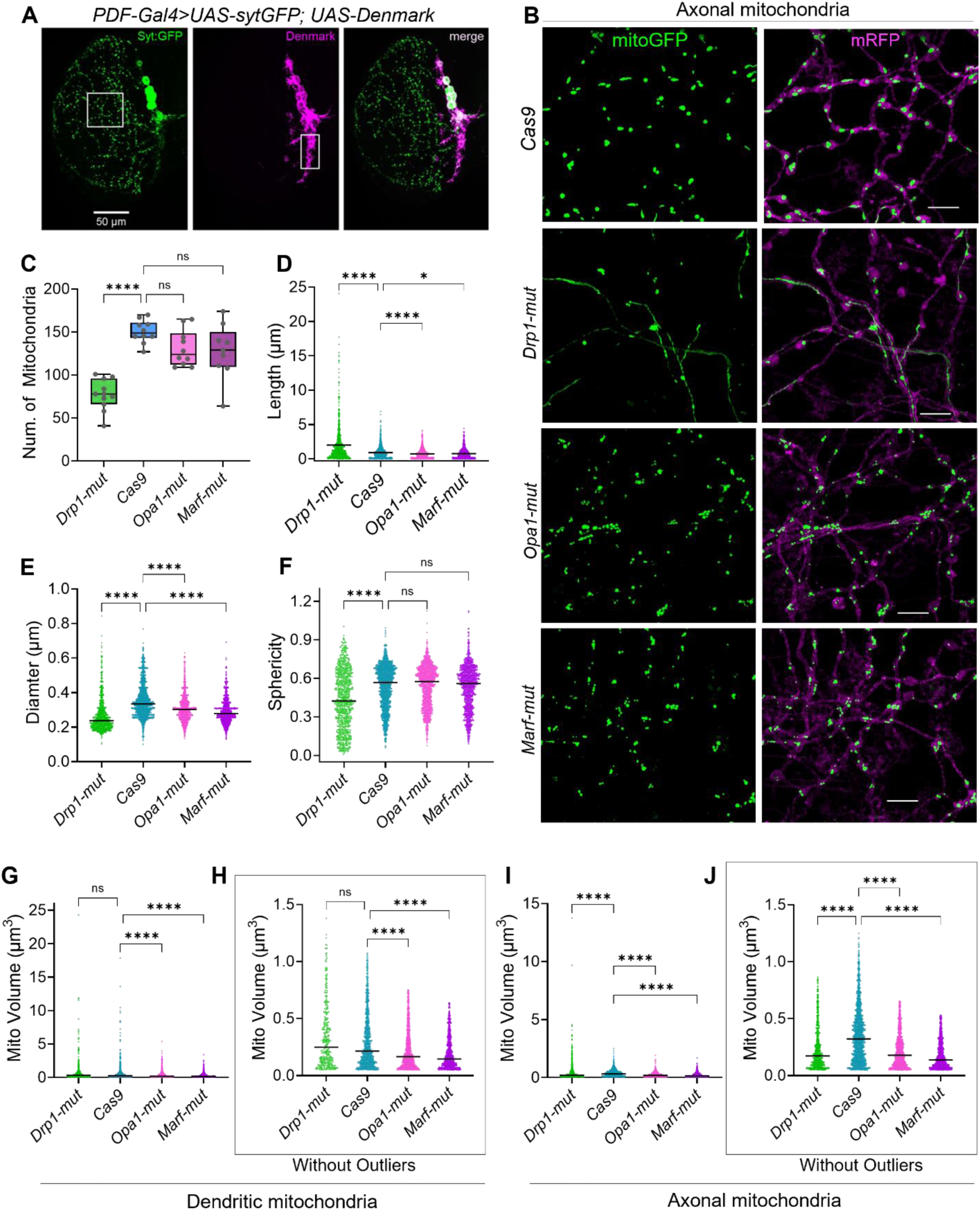
Cell type-specific CRISPR effectively clamps mitochondrial morphology. **A.** Dendritic and axonal projections of PDF neurons labeled with Denmark and SytGFP respectively. White boxes indicate approximate regions used to assay dendritic and axonal mitochondria **B**. Representative images of mitochondria in axons of the large-PDF neurons with indicated perturbations. Scale bars represent 5µm. **C.** Quantification of numbers of axonal mitochondria from the same approximate axonal region. *Drp1-mut* had much fewer axonal mitochondria than controls or fusion mutants, One-way ANOVA with Dunnett’s multiple comparisons test was used to compare Cas9 control and the mutants, adjusted p-value >0.05 is denoted as not significant (n.s.); ****p<0.0001. **D-F.** Length, diameter and sphericity of axonal mitochondria. **G-J.** Volume of dendritic and axonal mitochondria. For better visualization, H and J have outliers removed. Statistical significance is unchanged with or without outliers. Kruskal-Wallis ANOVA with a post hoc Dunn’s test was used to test statistical significance between Cas9-control and the mutants, adjusted p-value >0.05 is denoted as not significant (n.s.); *p<0.05; ****p<0.0001.

**Fig. S4:**
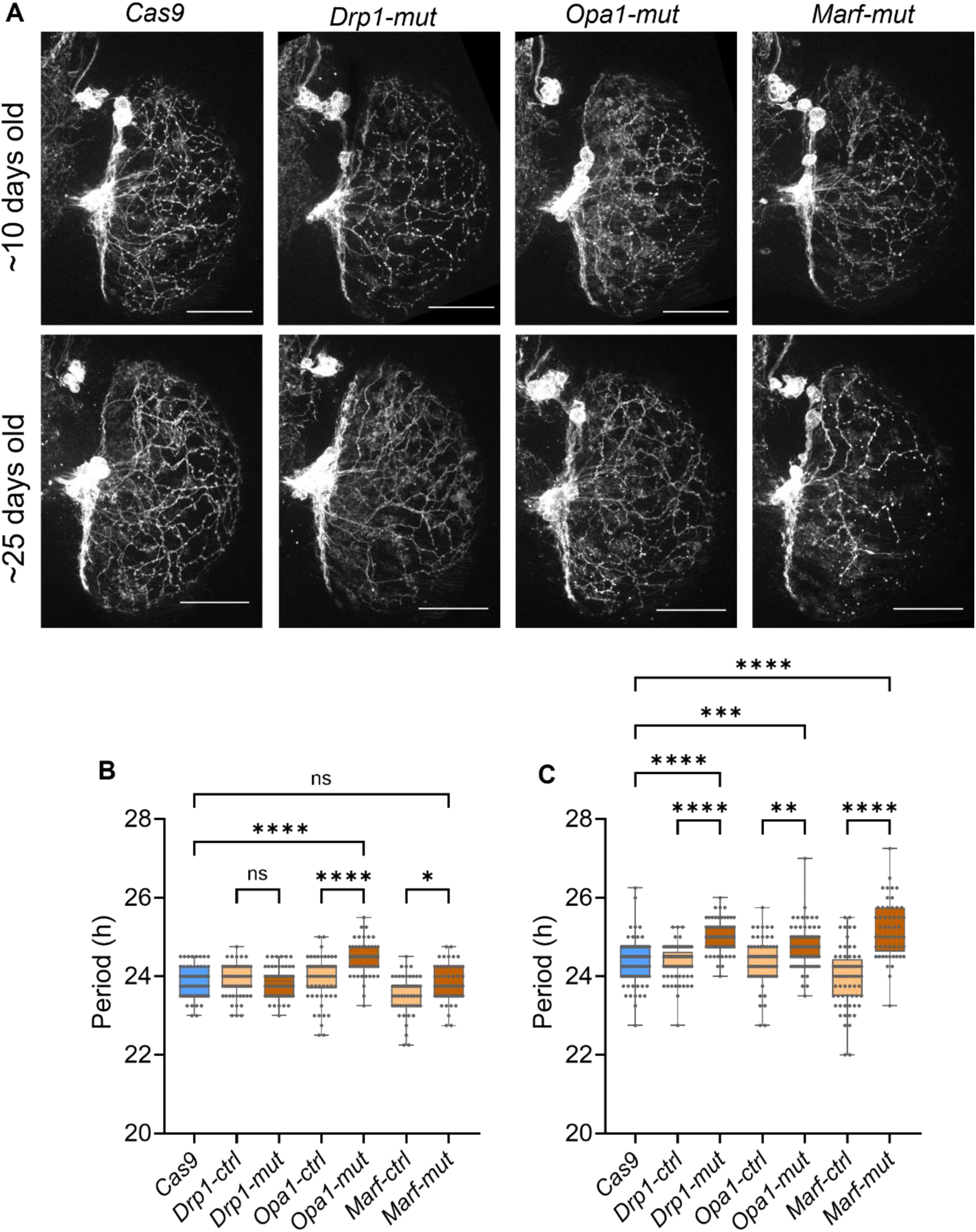
Loss of fission or fusion in old clock neurons leads to longer circadian periods. **A.** Representative images of ∼10 days and ∼25 days old ventral clock neurons and their projections labeled by *UAS-mRFP* expressed by *CLK856-Gal4* **B-C.** Circadian period of young (B, ∼10 days old) and old (C, ∼40 days old) flies of indicated genotypes. Period was calculated for DD days 2-7 for rhythmic flies (RI>0.3). Kruskal-Wallis ANOVA with a post hoc Dunn’s test was used to test statistical significance between mutants and Cas9-control as well as their respective gRNA controls, adjusted p-value >0.05 is denoted as not significant (n.s.); *p<0.05; **p<0.01; ***p<0.001; ****p<0.0001.

**Fig. S5:**
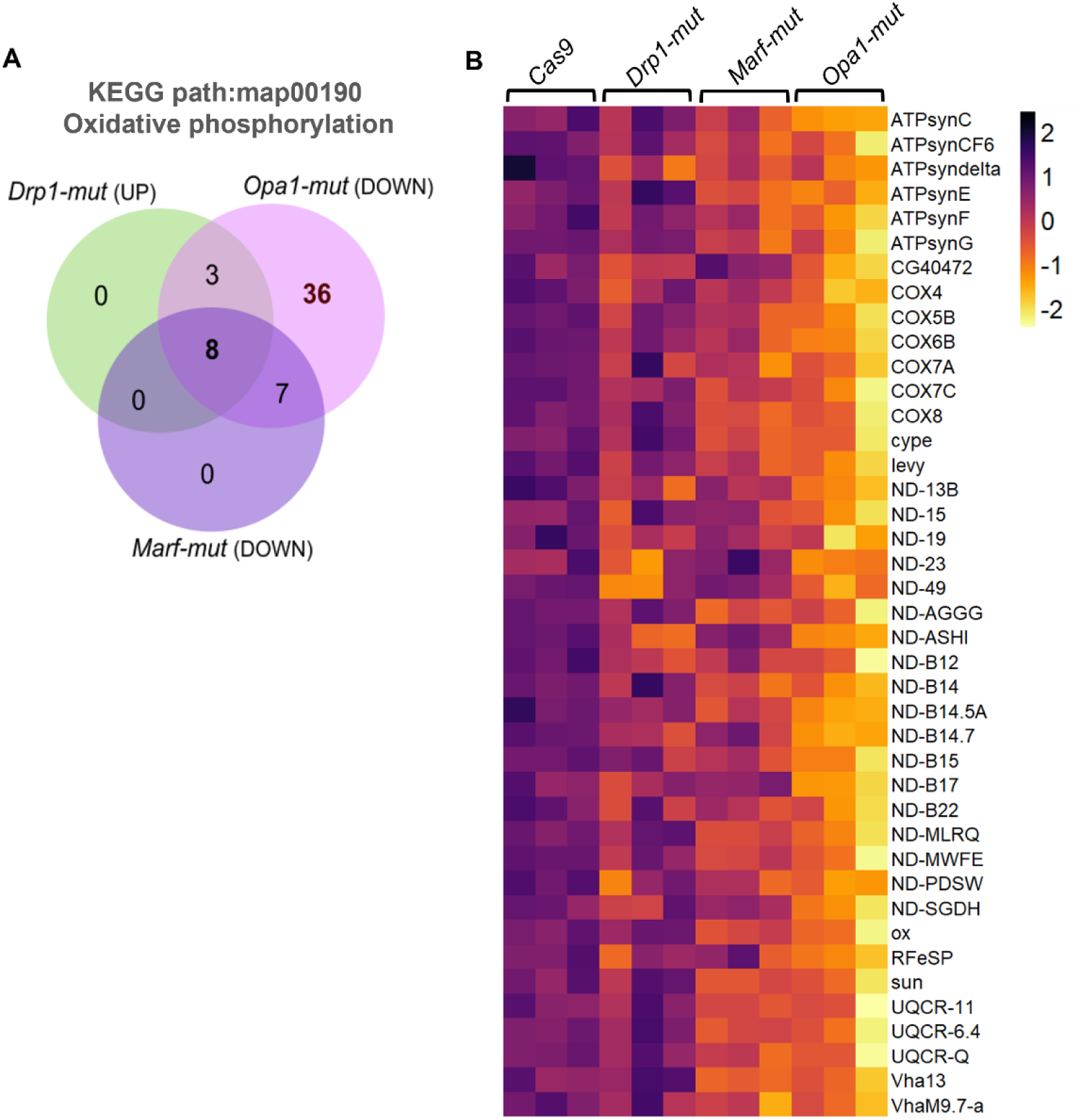
Loss of mitochondria fusion through *Opa1-mut* leads to downregulation of genes involved in oxidative phosphorylation. **A.** Venn-diagram of differentially expressed genes (upregulated in *Drp1-mut* and downregulated in both *Opa1-mut* and *Marf-mut*) belonging to the KEGG pathway category of oxidative phosphorylation in old clock neurons. *Opa1-mut* clock neurons is the superset, with most genes. All 8 genes common to the three conditions are encoded by mtDNA (See Fig. 3I). **B.** Heatmap of nuclear genes involved in oxidative phosphorylation downregulated in old *Opa1-mut* clock neurons.

**Fig. S6:**
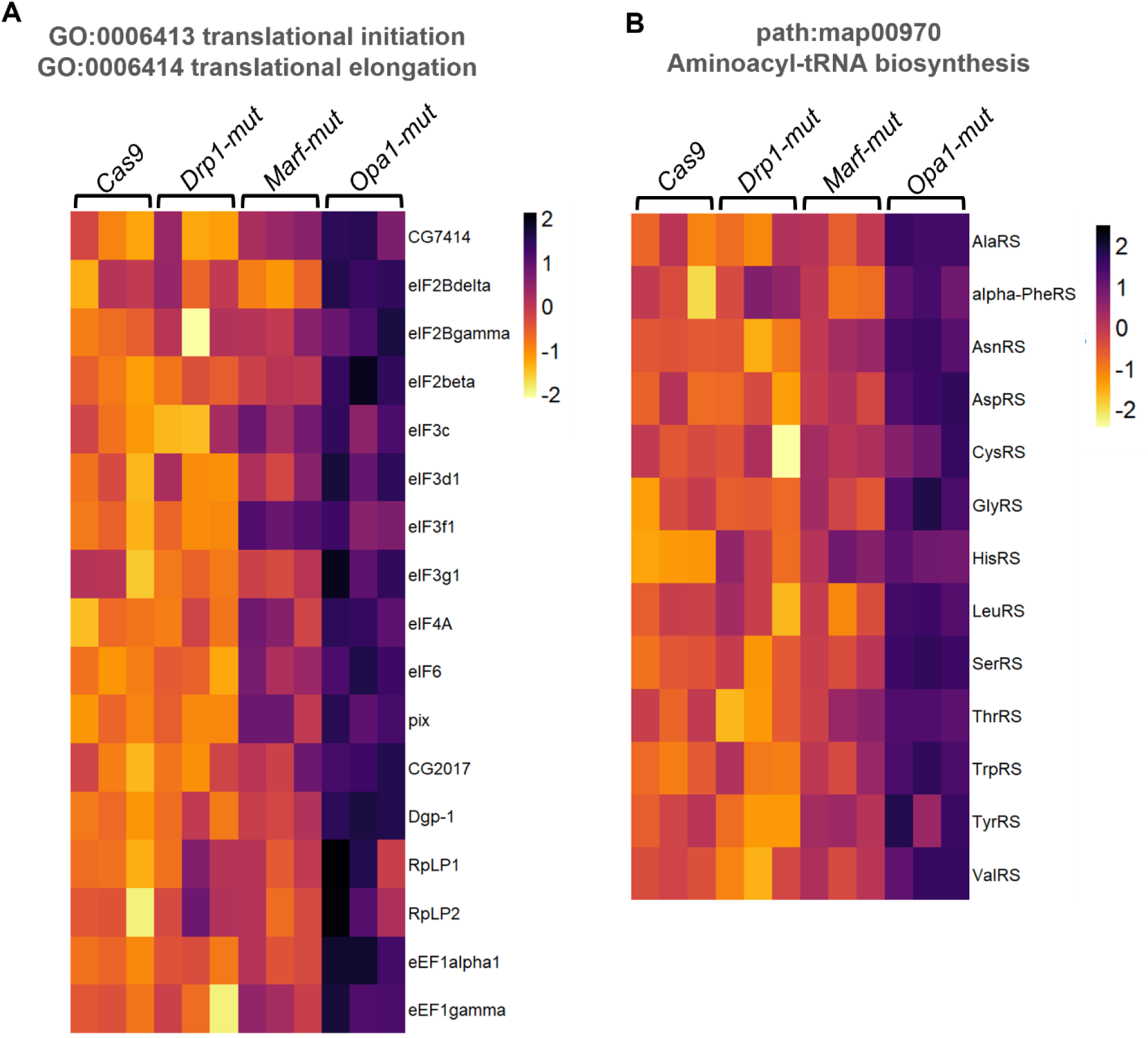
Loss of mitochondrial fusion through *Opa1-mut* leads to up-regulation of translation-related genes. Heatmaps of genes belonging to gene-ontology categories of translation initiation and elongation (**A**) and aminoacyl t-RNA biosynthesis (**B**) that were significantly up-regulated in old clock neurons.

**Fig. S7:**
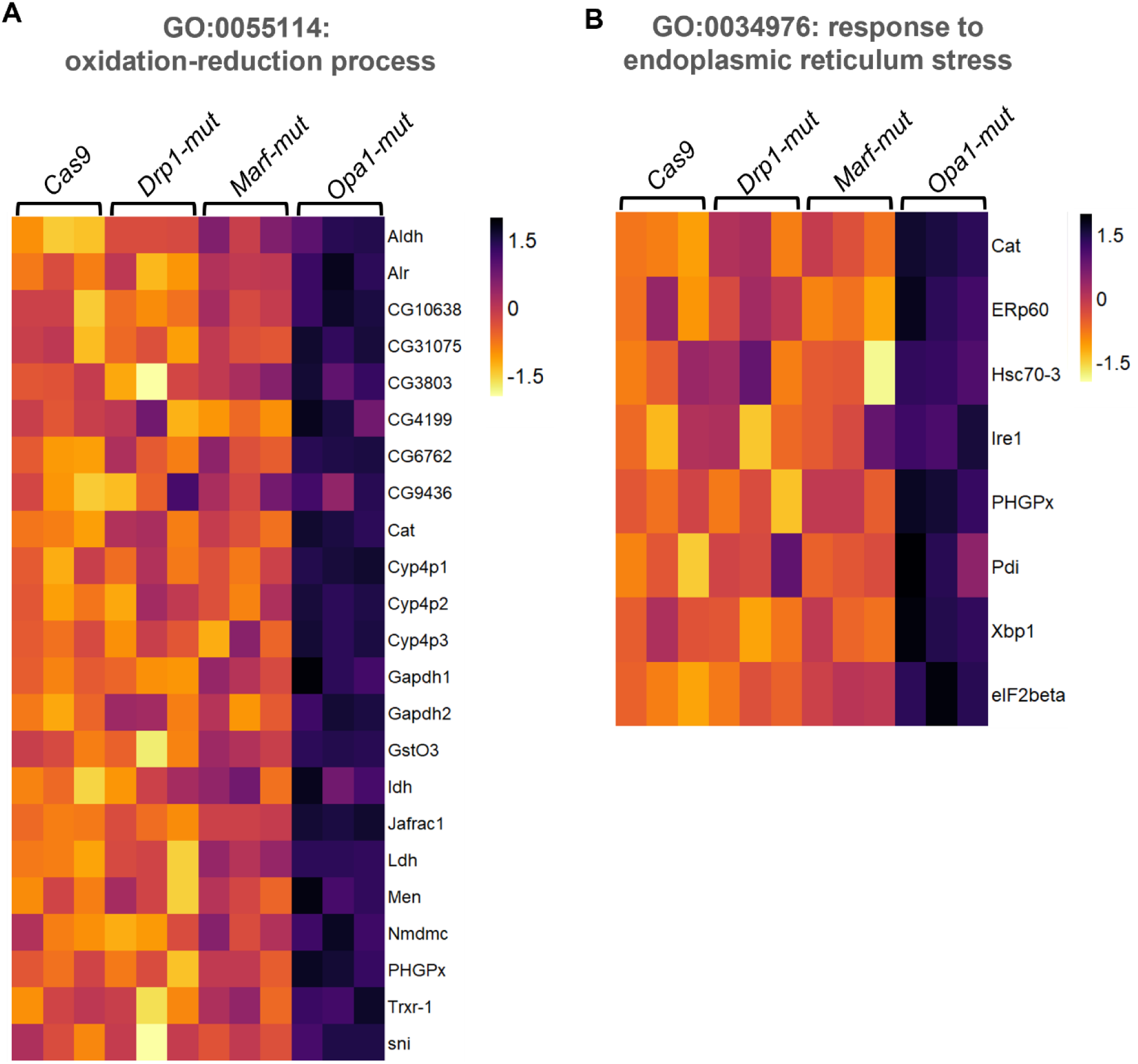
Loss of mitochondrial fusion through *Opa1-mut* leads to up-regulation of redox and ER stress genes. Heatmaps of genes belonging to gene-ontology categories of oxidation-reduction process (**A**) and response to endoplasmic reticulum stress (**B**) that were significantly up-regulated in old clock neurons.

**Fig. S8:**
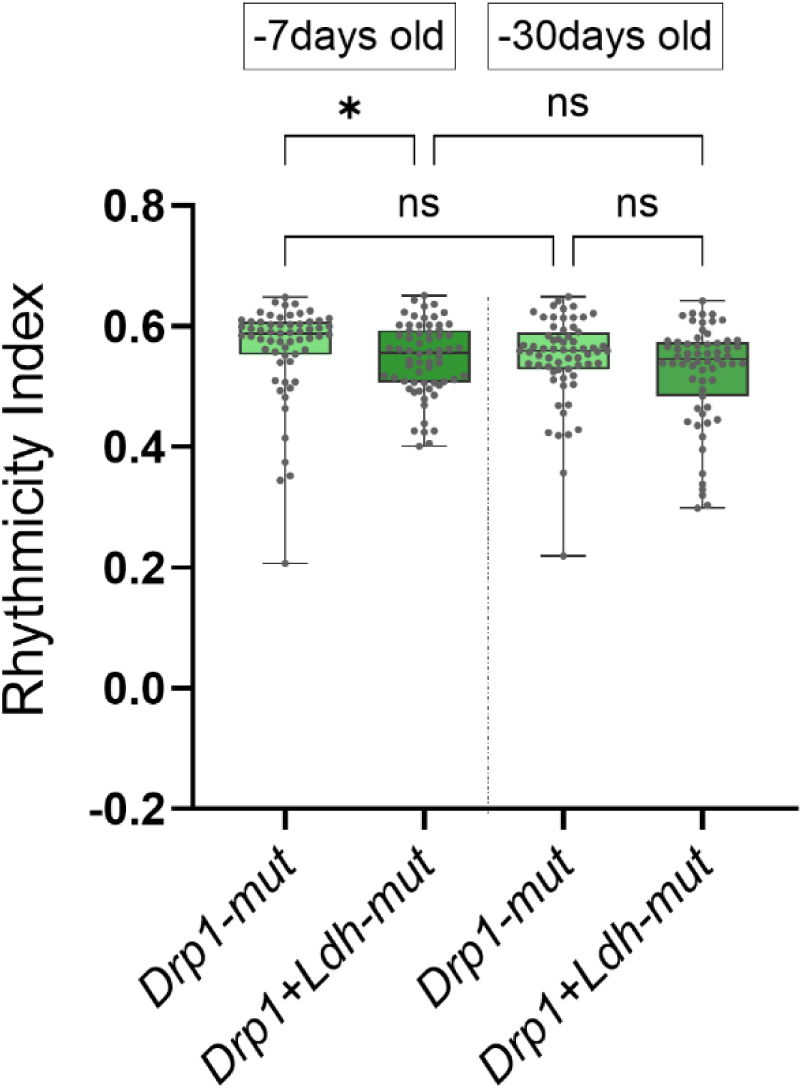
Loss of fission through *Drp1-mut* does not interact with *Ldh*. Rhythmicity of young (∼7 days old) and old (∼30 days old) with indicated mutations driven by *CLK856-Gal4* measured as described (Fig. 2 legend); n ≥ 60 flies per genotype from two independent experiments. Additional loss of *Ldh* in fission deficient clock neurons does not alter rhythmicity. Kruskal-Wallis ANOVA with a post hoc Dunn’s test, adjusted p-value >0.05 is denoted as not significant (n.s.); *p<0.05.

**Fig. S9:**
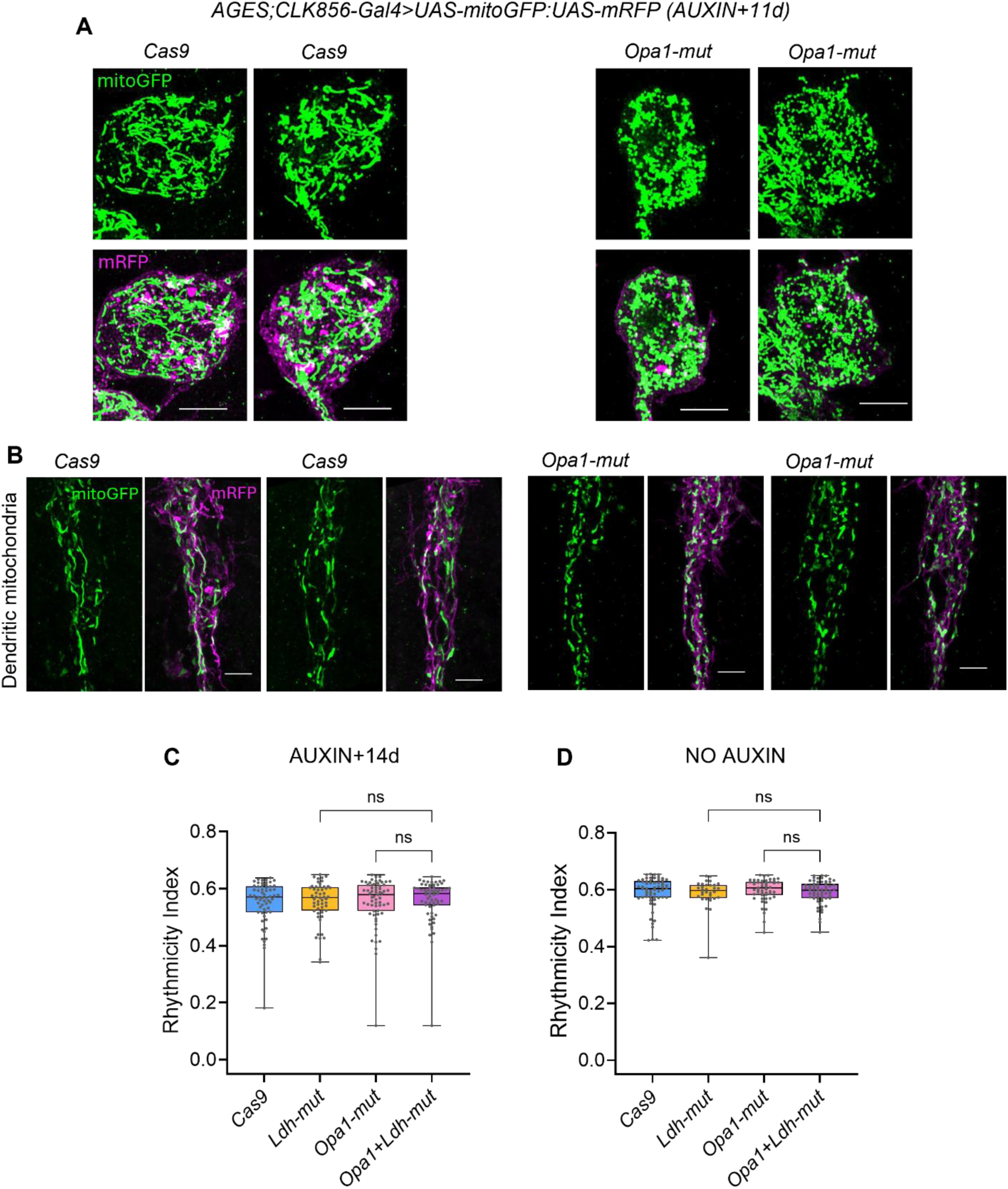
Chronic loss of mitochondrial fusion and *Ldh* is required for a functional effect. **A.** Two representative images each of large PDF neurons from indicated genotypes. *CLK856-Gal4* was combined with the adult-specific AGES system to induce mutations in the *Opa1* gene in clock neurons adult-specifically, as well as to visualize mitochondria using *UAS-mitoGFP* and label the neurons using *UAS-mRFP.* 2-3d old adult flies were fed on auxin containing food for 11 days, which induced fragmentation in mitochondrial morphology in most cells. **B.** Two representative images each of dendritic mitochondria from the same brains as described for (A). **C.** Rhythmicity Index of flies containing AGES, an auxin inducible expression system combined with *CLK856-Gal4* that were fed auxin for 5 days and tested after 14 days from the first day of auxin feeding, n ≥ 53 flies for each genotype from at least two independent experiments **D.** Rhythmicity Index for flies of the same genotypes as (C) aged to ∼30d but never fed auxin, n ≥ 32. *Opa1+Ldh-mut* flies, either tested 14 days post auxin or old flies without any auxin had no loss of rhythmicity, Kruskal-Wallis ANOVA with a post hoc Dunn’s test, adjusted p-value >0.05 is denoted as not significant (n.s.). This indicates a chronic loss of *Opa1+Ldh* is required for loss of function (Also see Fig. 5D-F).

**Fig. S10:**
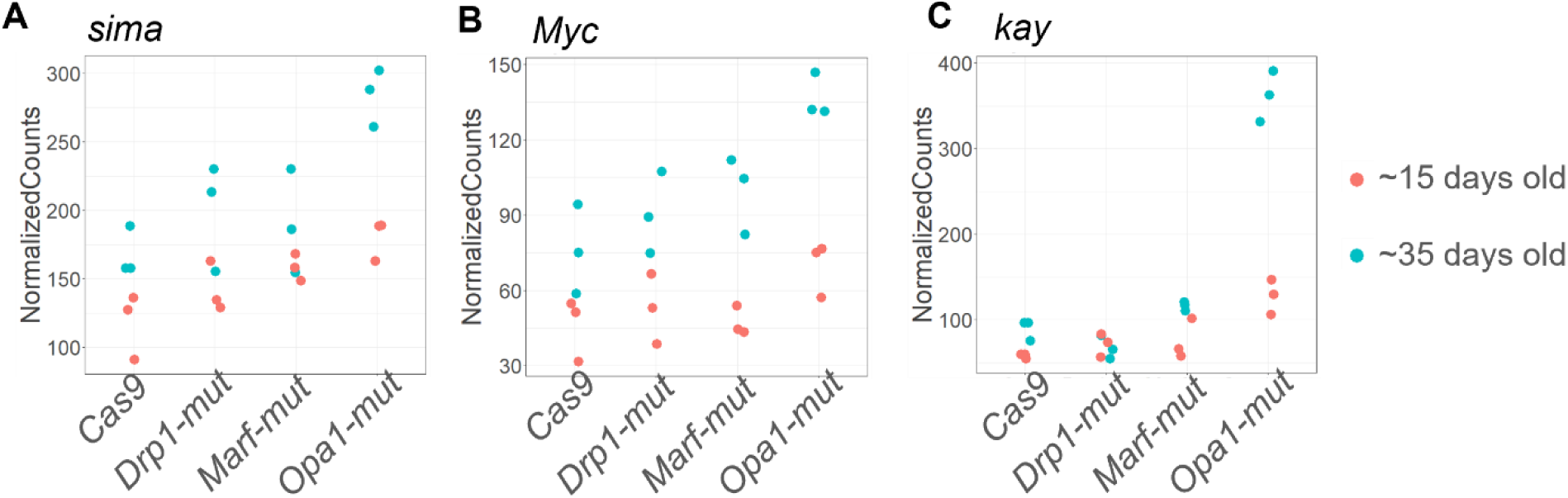
Transcription factors associated with the Warburg effect are up-regulated in *Opa1-mut* old clock neurons. Normalized counts for *sima, Myc* and *kay* from the bulk sequencing dataset.

## Supplementary Tables

**Table S1:** List of fly stocks.

| Genotype | Description | Source/Reference |
| --- | --- | --- |
| *CLK856-Gal4* | GAL4 labelling most clock neurons | (Gummadova et al., 2009) |
| *PDF-Gal4* | GAL4 labelling the PDF-positive clock neurons | (Renn et al., 1999) |
| *UAS-mitoGFP* | Express GFP targeted to the mitochondria under UAS-control | BDSC 8443 |
| *UAS-SytGFP:Denmark* | Express SytGFP (labeling axons) and -Denmark (labeling dendrites and cell bodies) under UAS-control | BDSC 33065 |
| *UAS-mRFP* | Express mCD8-RFP under UAS-control | BDSC 32218 |
| *UAS-eGFP* | Express eGFP under UAS-control | BDSC 5430 |
| *UAS-Cas9 (II)* | Express Cas9.P2 under UAS-control (II chromosome) | BDSC 58985 |
| *UAS-Cas9 (III)* | Express Cas9.P2 under UAS-control (III chromosome) | BDSC 58986 |
| *UAS-Drp1-g* | Express 3x-gRNA line against Drp1 under UAS-control | this study |
| *UAS-Opa1-g* | Express 3x-gRNA line against Opa1 under UAS-control | this study |
| *UAS-Marf-g* | Express 3x-gRNA line against Marf under UAS-control | this study |
| *UAS-Ldh-g* | Express 3x-gRNA line against Ldh under UAS-control | this study |
| *UAS-ATF4-g* | Express 2x-gRNA line against ATF4 under UAS-control | VDRC 341284 |

**Table S2:**
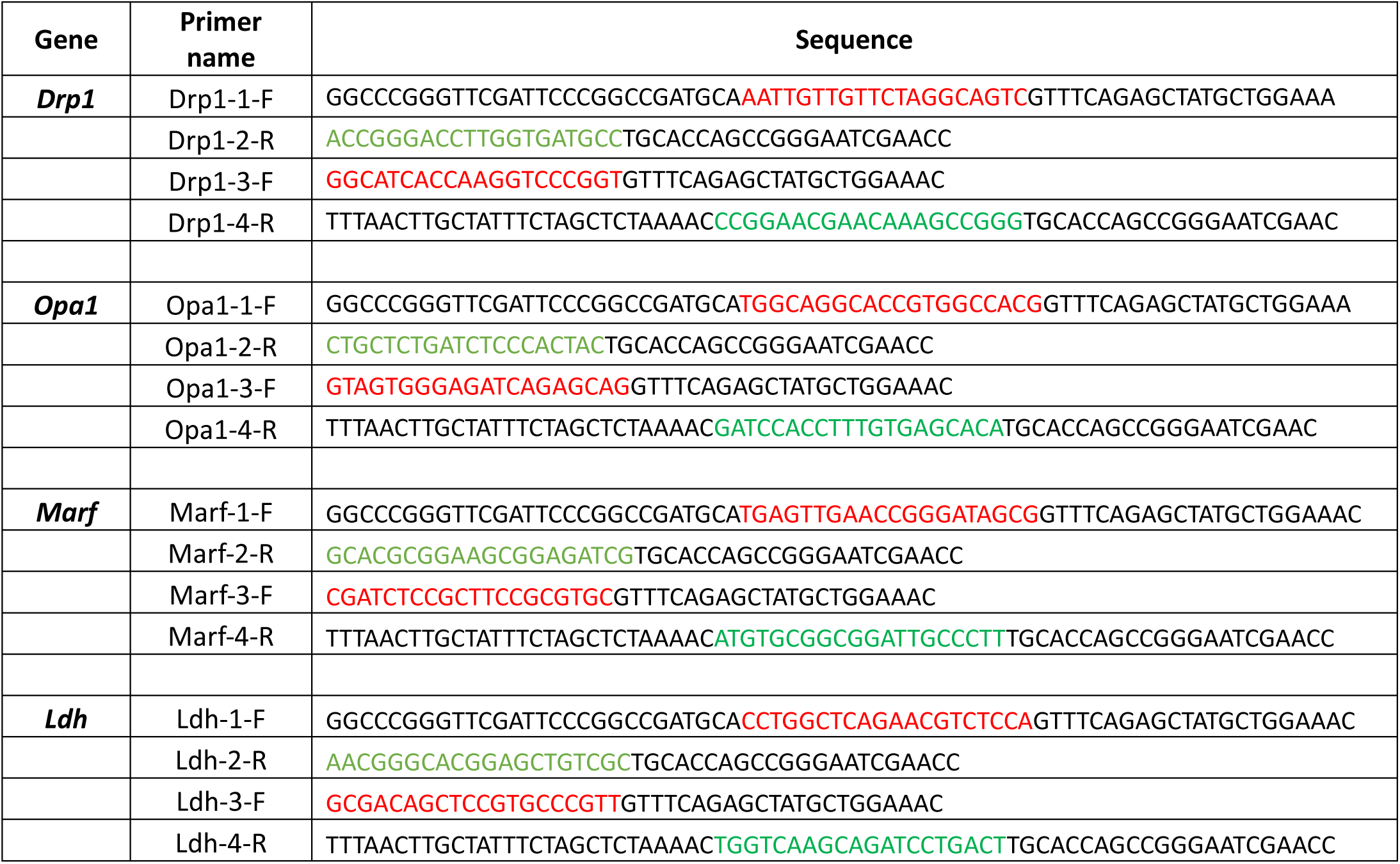
Primer sequences used for cloning gRNA lines. Colored portions are gRNA sequences.

